# Characterizing features of Kv channel activation currents by a minimal-state kinetic model

**DOI:** 10.64898/2026.10.26.761847

**Authors:** Di Wu, Shushang Jiang, Yiyun Huang

## Abstract

Voltage-gated Potassium channels (Kv channels) comprise twelve subfamilies found in the human genome. They demonstrate diversified activation behaviors, represented by the varied shapes of activation currents, which are usually modeled by individual kinetic schemes of many parameters that make channel comparisons difficult. Here, we present a minimal-state kinetic model, aiming to compare different Kv channel activation currents by a unified model system. By omitting the short-lived transient states and neglecting the detailed transition pathways, this model employs only minimal significant states necessary to approximate the activation currents. Few model-parameters are used that capture the varied current-features, leading to a convenient comparison among Kv channels. In addition, this model incorporates a gating factor, which accurately represents the channel opening event upon applying a voltage pulse. A special feature of the model is its adaptability via simple modifications, that any channel-dependent kinetic scheme can be easily incorporated into this model framework, helping develop exact channel mechanisms.

## Introduction

In 1952, Hodgkin and Huxley published their famous kinetic model (abbreviated to HH model here), characterizing sodium- and potassium-channel activation currents that successfully simulated nerve impulse propagations observed in the squid giant axons^1^. Following their seminal work, more ion-channel models were developed, encompassing a variety of Markov-type models and parameter-estimation methods based on single-channel properties^2–24^. These models are very helpful in describing diversified ion-channel behaviors and they are often used in neuronal and cardiac electrophysiological simulations. In addition, other types of models and modeling methods are also developed^25–31^, and models based on artificial-intelligence methods are emerging recently^32–35^, further improving ion-channel modeling procedures.

Despite these achievements, the issue of model identifiability remains^27,29,36,37^ ever since the development of ion-channel models especially the Markov-type models, that a specific channel behavior can be represented by several model schemes or layouts all with acceptable results. Although methods were developed that helped select or construct the optimum model scheme^8,12,27,29,30^, the procedure of model selection is still difficult. Currently, thousands of ion-channel models are archived at ModelDB^38,39^ website (https://modeldb.science). This further complicates model selections if one has no idea of a proper model layout initially. In 2017, Podlaski *et al.* had successfully reduced 2378 voltage- and calcium-gated ion channel models (coded in NEURON^40^ available at ModelDB website) into 211 clusters^41^, greatly facilitating the model-selection procedure. Apart from selecting models from established schemes, here, we try to construct a unified model scheme suitable to represent (approximately) many ion-channel behaviors, thus circumventing the model-selection step while enabling cross-channel comparisons. Such a model system can be practicable only for a group of ion channels demonstrating common conduction behaviors, and Kv channel family may be suitable for the study.

Voltage-gated potassium channels comprise 12 subfamilies found in the human genome^42^. They share common features in their structures^43,44^, including six transmembrane helices in each α subunit, where the voltage-sensor (helices S1-S4) motions^45–48^ elicited by a voltage change can trigger the opening or closing of the gate located at the pore region (helices S5-S6 including a pore-loop structure) that conducts ionic currents. Kv channels show varied inactivation behaviors^49–51^, including N-type^50^ and C-type inactivations^52–59^. Their delayed response to depolarizing voltages and varied inactivation behaviors greatly diversify their activation currents, making them suitable for modeling studies.

The work of unifying mathematical models to fit ion-channel currents emerges recently, showing promising results^35,60^. Linkevicius *et al.* proposed a machine-learning method combined with nonlinear-mixed-effects modeling based on Hodgkin-Huxley model system that successfully modeled potassium currents of 20 Kv channel subtypes^35^. Their work greatly improves Kv channel model-fitting procedures based on the HH model system. Besides this, a unified kinetic scheme implemented in Markov model was proposed by Balbi *et al.*, who constructed a six-state kinetic scheme that successfully modeled Na_v_1.1 to Na_v_1.9 channel currents^60^.

Different from their work, here, we present a unified and improved Markov-type model framework, which is adaptable that can employ varied kinetic schemes, encompassing not only the unified kinetic scheme for fitting most Kv channels (which is the purpose of this study), but also other schemes describing more detailed mechanisms for individual channel behaviors (not examined in this study). Thus, this model framework enables researchers to revise (conveniently) a kinetic scheme based on an initial guess (e.g., the unified kinetic scheme). The advantage of using this framework is the employment of fewer kinetic states that both fit current well and yield plausible mechanisms inferred from the model-parameter values.

We check this model using experimental data obtained from Channelpedia^61^ website (https://channelpedia.epfl.ch), where Ranjan *et al.* have comprehensively studied and archived various Kv channel currents^62^. Their work greatly enriches our knowledge of different Kv channel behaviors, making Kv channel comparisons much easier. From their website, we have selected 54 Kv channel activation currents comprised of 18 Kv channel subtypes recorded at three temperatures. Among them, 41 currents were fitted well by a four-state kinetic scheme, and the others were fitted by a five-state kinetic scheme. Thus, the four-state kinetic scheme may be used as a unified model scheme for fitting Kv channel activation currents of unknown mechanisms initially, or for comparing cross-Kv channel activation behaviors approximately.

The viability of the unified model scheme relies on the common conduction behavior of Kv channels, which may occur in their activation currents lasted for a short time. Here, we do not model Kv channel deactivation, inactivation, or recovery currents, which may require additional states or state-transitions that can complicate the kinetic scheme. Instead, we focus on fitting only activation currents, where similar states may occur chronologically over a short period. By incorporating a gating factor and utilizing a voltage-dependent current level for each state (excluding the first state), our model can fit different Kv channel activation currents at high-level accuracies (depending on the number of states employed). The results are presented and discussed in the following sections.

## Model

We present a minimal-state kinetic model (abbreviated to MS model) to capture the main features of Kv channel activation currents over 500 ms time period. We aim to find a minimal number of states suitable to describe various Kv channel activation behaviors via a pseudo-linear kinetic propagation scheme. Our experiments with Channelpedia data^62^ show that four states implemented in the MS-model framework are generally enough to characterize most Kv channel activation currents, although some channels do require five or more states for more accurate representations.

Normally, a kinetic model describing a channel behavior can be complex, comprising many microscopic states connected by a network of state-transitions. Among these states, several are usually employed to represent the closed-channel conditions, which are consistent with the experimental observations^63^. Here, we have concatenated all closed states into one non-conducting State *A*. Furthermore, any branched or cyclic transitions normally occurred in a channel mechanism are truncated, leaving only a linearly forward propagation. This means that all Kv channels are forced to follow the same pseudo kinetic scheme, which may not be an exact mechanism for individual channels. To develop channel-dependent mechanisms, further scheme-modifications are needed, that can be conveniently implemented under the same MS-model framework. Thus, depending on individual needs, researchers may choose different kinetic schemes for their investigations.

The MS-model schemes we use in this study are the four-state (Eq. 1) and five-state (Eq. 2) linear-propagation schemes, employing forward rate constants only. These pseudo-linear kinetic schemes are feasible to approximate the activation currents when the forward rate constants are much larger than the reverse rate constants. In the following derivations, kinetic parameters are expressed as a function of voltage only. Other environmental factors, such as temperature, can be included in the expression in the same way.

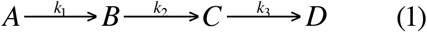

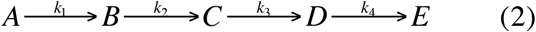

Here, *k_j_* (*j* = 1∼4) denotes rate constant. *A*, *B*, *C*, *D*, and *E* denote different states (commonly represented by *S_j_*, *j* = 1∼5), and their populations are calculated by Eq. 3, where only voltage-dependence is included in the parameter expressions:

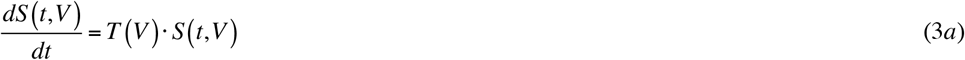

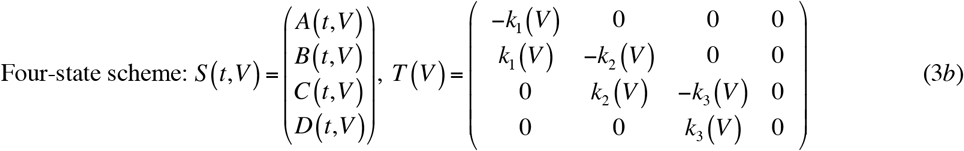

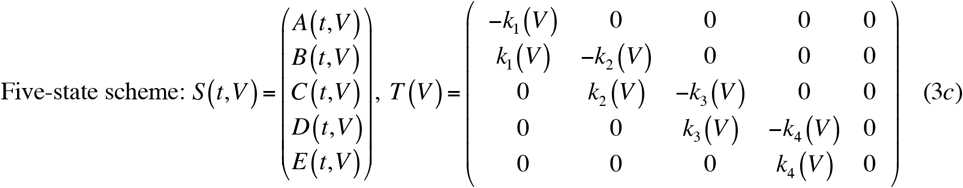

Here *t* denotes time, *V* denotes voltage, *S* denotes state ensemble, and *T* denotes state-transition matrix. For deriving other kinetic schemes, more states can be included by modifying *S*, and different state-transitions can be included by modifying the transition matrix *T*.

The beginning period of the activation current is usually difficult to model. We notice that, for Kv channels, the opening (triggered by the sliding motions of voltage-sensors^45–48^) of the voltage-sensor controlled gate is time-dependent. The voltage-sensors move at a fast speed preceding the permeation of ions through Kv channel’s pore region. Here, this time-dependent opening of the voltage-sensor controlled gate is incorporated into the MS model, which is represented by a gating factor *GF*:

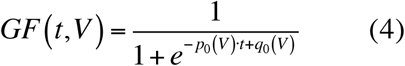

Here *t* denotes time, *V* denotes voltage, *p*_0_ and *q*_0_ are model parameters. The role of this gating factor is to represent a situation where the ionic current does not occur unless the voltage-sensor controlled gate is open. With this gating factor, the results are more accurately represented at the beginning period of the activation current.

Due to the minimal states employed, each state may be a composite of several substates of same or similar subconductance levels. This may change the state-current (irregularly in some cases) over a range of voltages. Therefore, each state (except State *A*) of MS model is assigned a parameter *i_j_* to represent the reduced current level associated with it. This design helps in merging states and improves the fitting qualities while forcing all channels to follow the same linear and forward-only kinetic propagations.

In this way, the framework of macroscopic current represented by the MS model is:

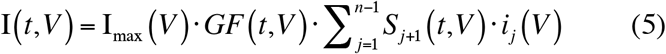

Here I_max_ denotes the maximum current at a given voltage, which becomes a model parameter that is conveniently obtained by finding the maximum value on a given current curve. The gating factor *GF* is calculated by Eq. 4 and the state-population *S_j_* is calculated by Eq. 3. *n* denotes the number of states employed, *n* = 4 or 5 for the four- or five-state scheme, respectively.

In this study, 9 parameters are used for the four-state scheme (I_max_, *p*_0_, *q*_0_, *k*_1_, *k*_2_, *k*_3_, *i*_1_, *i*_2_, and *i*_3_) and 11 parameters are used for the five-state scheme (I_max_, *p*_0_, *q*_0_, *k*_1_, *k*_2_, *k*_3_, *k*_4_, *i*_1_, *i*_2_, *i*_3_, and *i*_4_). All parameters are non-negative. *i_j_* value is normally between 0 and 1. It can be larger than 1 if I_max_ occurs at a time when *S_j_* values are small, which sometimes can result from the uncertainties due to the small-scale currents recorded at low voltages.

Kinetic model parameters can depend on many environmental factors including voltage. The rate constant *k_j_*in previous kinetic models usually followed an exponential voltage-dependence^64,65^. Here, with the inclusion of the gating factor, the rate constants are found linearly dependent on voltage approximately. Parameter *i_j_* can follow linear or exponential voltage-dependence, and their values can help describe the state properties. These will be discussed in the Results section.

Due to the stochastic nature of the activation current, priority is given to fitting each current instead of fitting the parameters to follow a voltage-dependent function. Results presented in this way help examine the viability of the MS model and prove that the model can fit currents well using limited states. On the other hand, the parameter-voltage plots are presented in Supplementary Figures S1-S18, where each parameter’s voltage-dependence can be examined and interpreted by a user-defined function.

## Results

### Fitting Kv channel activation curves by MS model

The activation currents over 500 ms time period vary greatly among Kv channel subfamilies, which are clearly shown in the experimental recordings^62^ archived at Channelpedia website (https://channelpedia.epfl.ch). For example, over 500 ms period, Kv10.1 channels show non-inactivating behaviors at low temperatures, Kv4 channels show fast-inactivating behaviors, Kv3.4 channels show various inactivating rates, whereas Kv5, Kv6, Kv8, and Kv9 channels are normally considered silent. In addition, some Kv channels, e.g., Kv1.8 channels, show small-scale currents. To avoid ambiguity in fitting currents, in this study, we have chosen only eighteen obviously conducting Kv channel subtypes, including Kv1.1, Kv1.2, Kv1.3, Kv1.4, Kv1.5, Kv1.6, Kv1.7, Kv2.1, Kv2.2, Kv3.1, Kv3.2, Kv3.3, Kv3.4, Kv4.1, Kv4.2, Kv4.3, Kv10.1, and Kv12.1 channels. Altogether, 54 Kv channel cell-current recordings are selected, comprising currents from 18 channel subtypes recorded at three temperatures (15 °C, 25 °C, and 35 °C). They demonstrate diversified activation behaviors, suitable for cross-channel comparisons.

We neglected fitting currents recorded below -20 mV, because large uncertainties existed due to either a non-opening channel or a tiny current-scale not separable from the background currents. Thus, the activation currents recorded over only -20 mV to 80 mV voltage-range, denoted as *n*8 to *n*18 curves, were fitted by the MS model. The results are shown in Figs. 1-5, and the model-parameter values are plotted in Supplementary Figures S1-S18, where the parameter-voltage relations can be examined. Generally, parameters I_max_, *p*_0_, *q*_0_, *k*_1_, *k*_2_, *k*_3_, and *k*_4_ show linear dependence on voltage, while *i*_1_, *i*_2_, *i*_3_, and *i*_4_ can be linearly or exponentially dependent on voltage. These will be discussed later.

**Figure 1.**
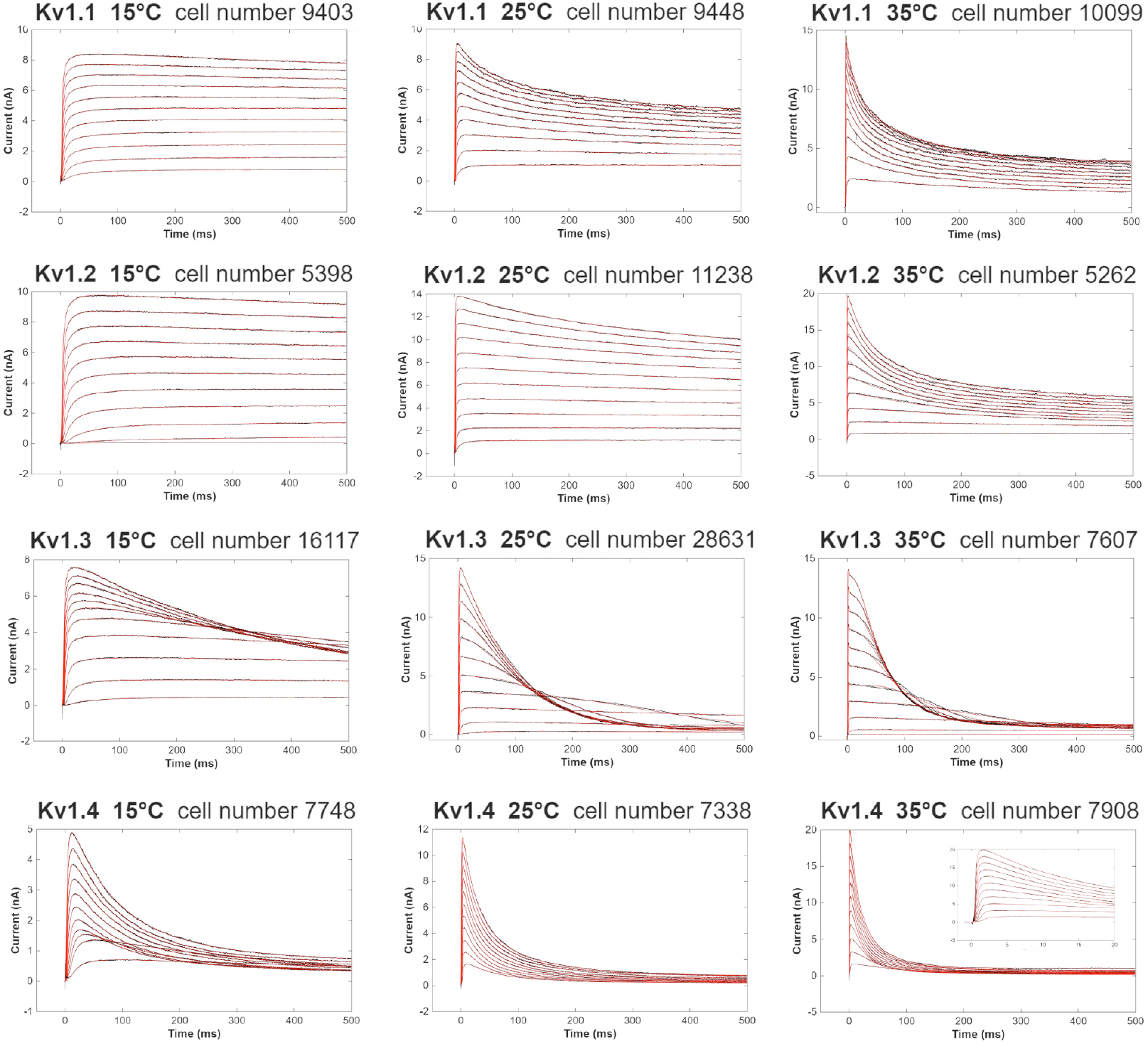
Kv channel activation currents fitted by the MS model. Eleven current traces (black curves, stimulated by voltages from -20 mV to 80 mV at 10 mV intervals) of each Kv (Kv1.1, Kv1.2, Kv1.3, and Kv1.4) channel recorded at three temperatures were fitted by the MS model (red curves). Each recording is labeled by a unique cell ID number obtained from Channelpedia website. The inset of Kv1.4 (35 °C) subplot shows the enlarged picture of the fitted currents during the initial 20 ms time period.

**Figure 2.**
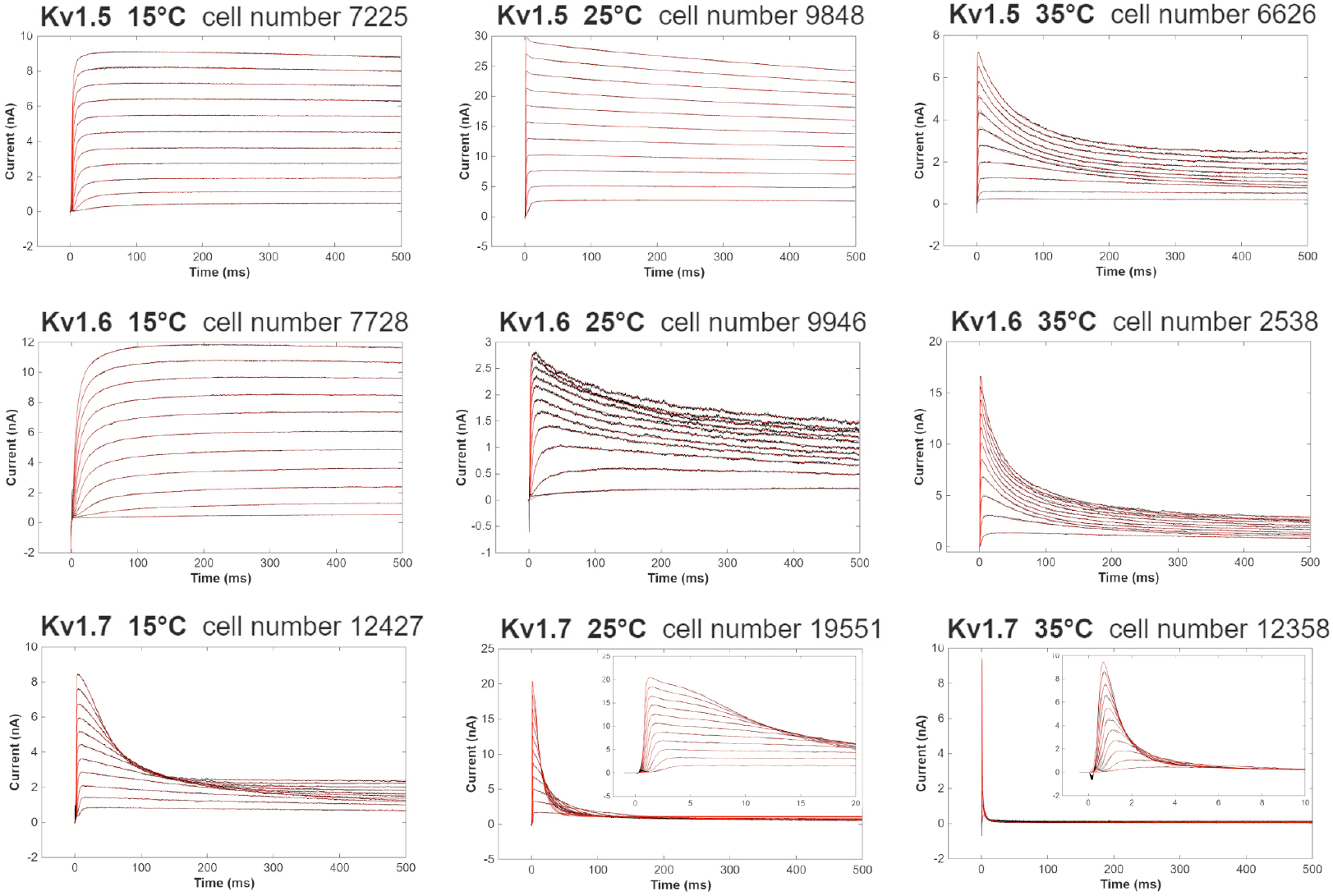
Kv1.5, Kv1.6, and Kv1.7 activation currents (black curves) fitted by the MS model (red curves). The insets show the enlarged pictures of the fitted currents at early time.

**Figure 3.**
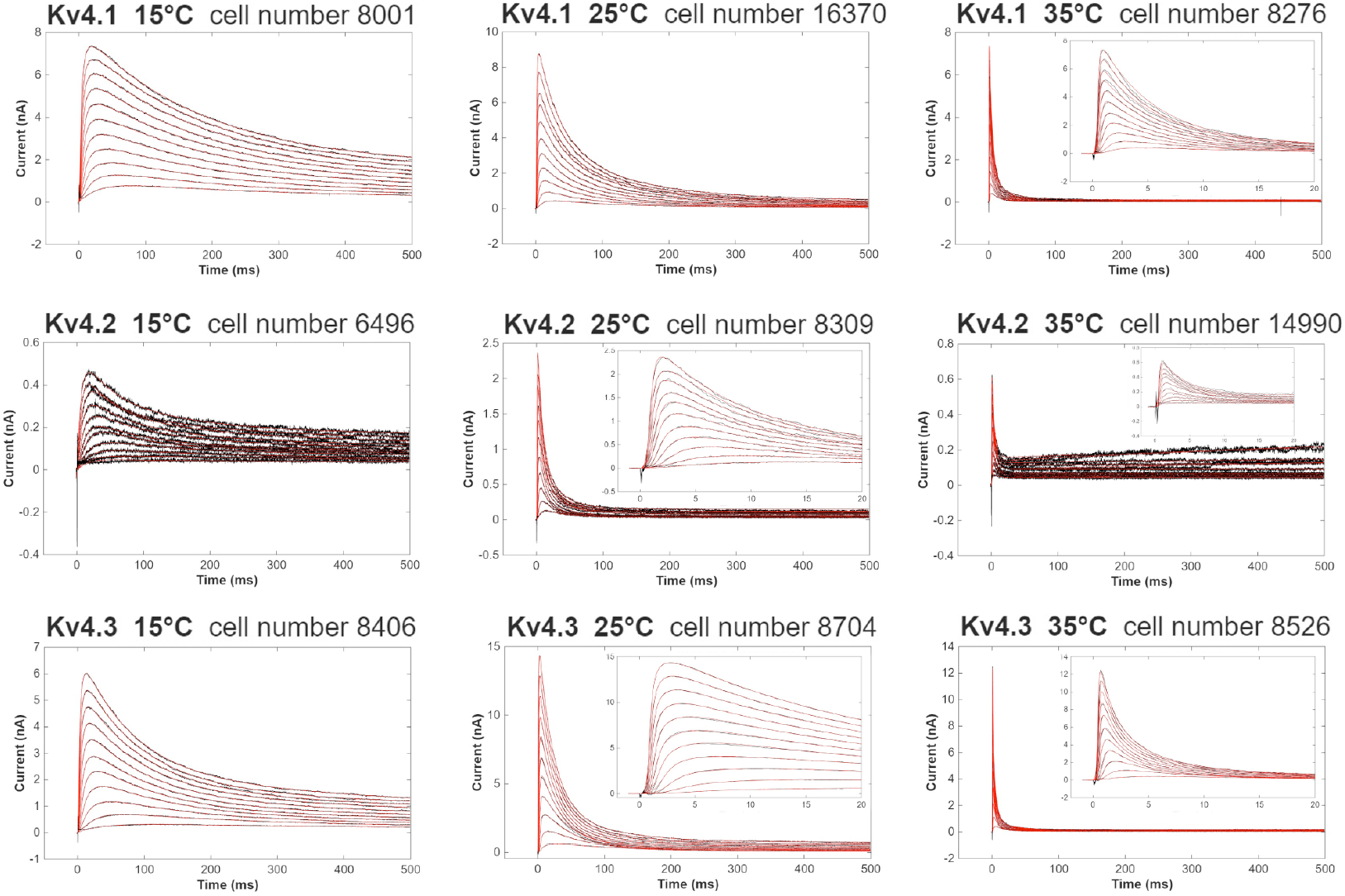
Kv4.1, Kv4.2, and Kv4.3 activation currents (black curves) fitted by the MS model (red curves). The insets show the enlarged pictures of the fitted currents at early time.

**Figure 4.**
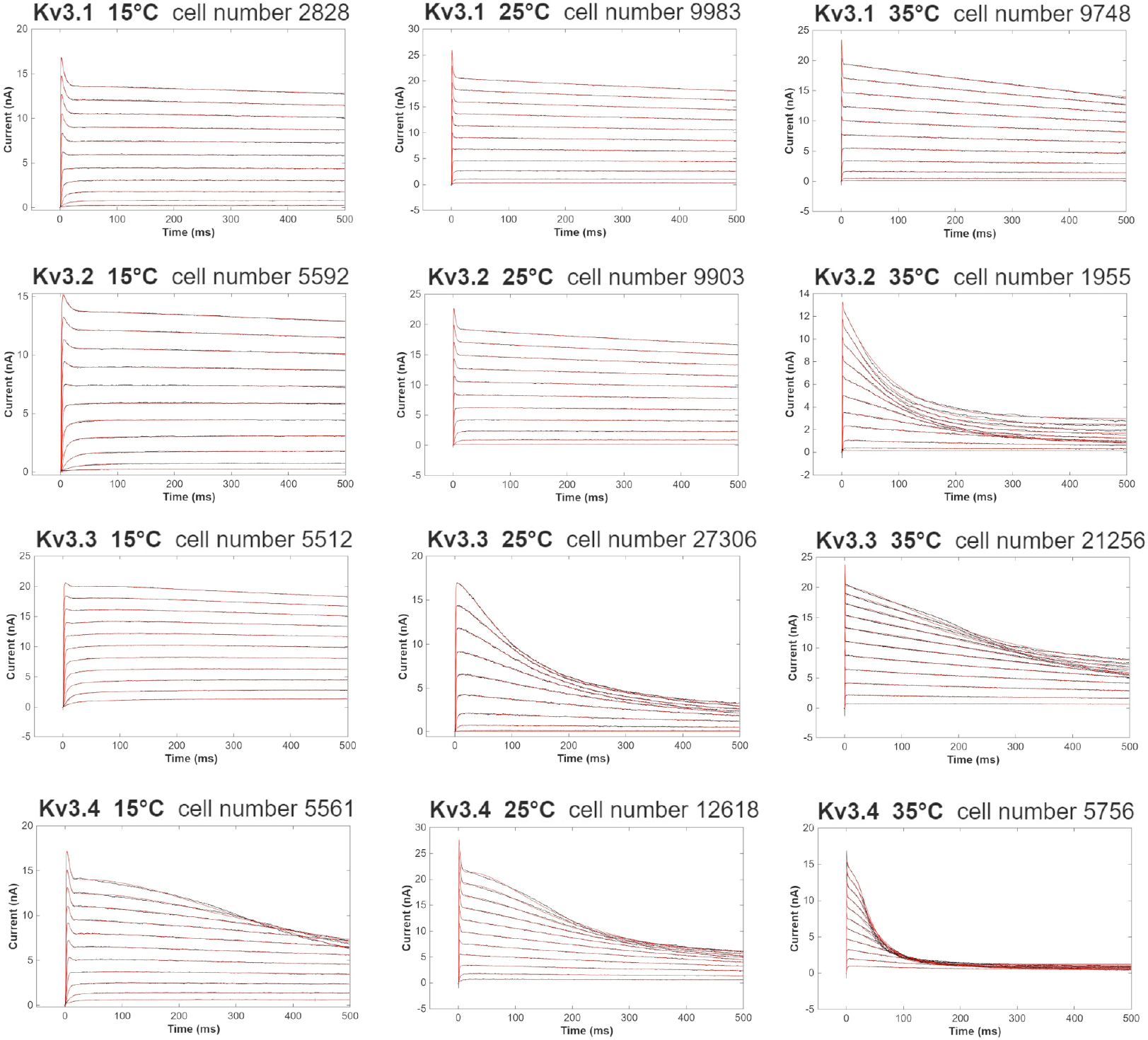
Kv3.1, Kv3.2, Kv3.3, and Kv3.4 activation currents (black curves) fitted by the MS model (red curves).

**Figure 5.**
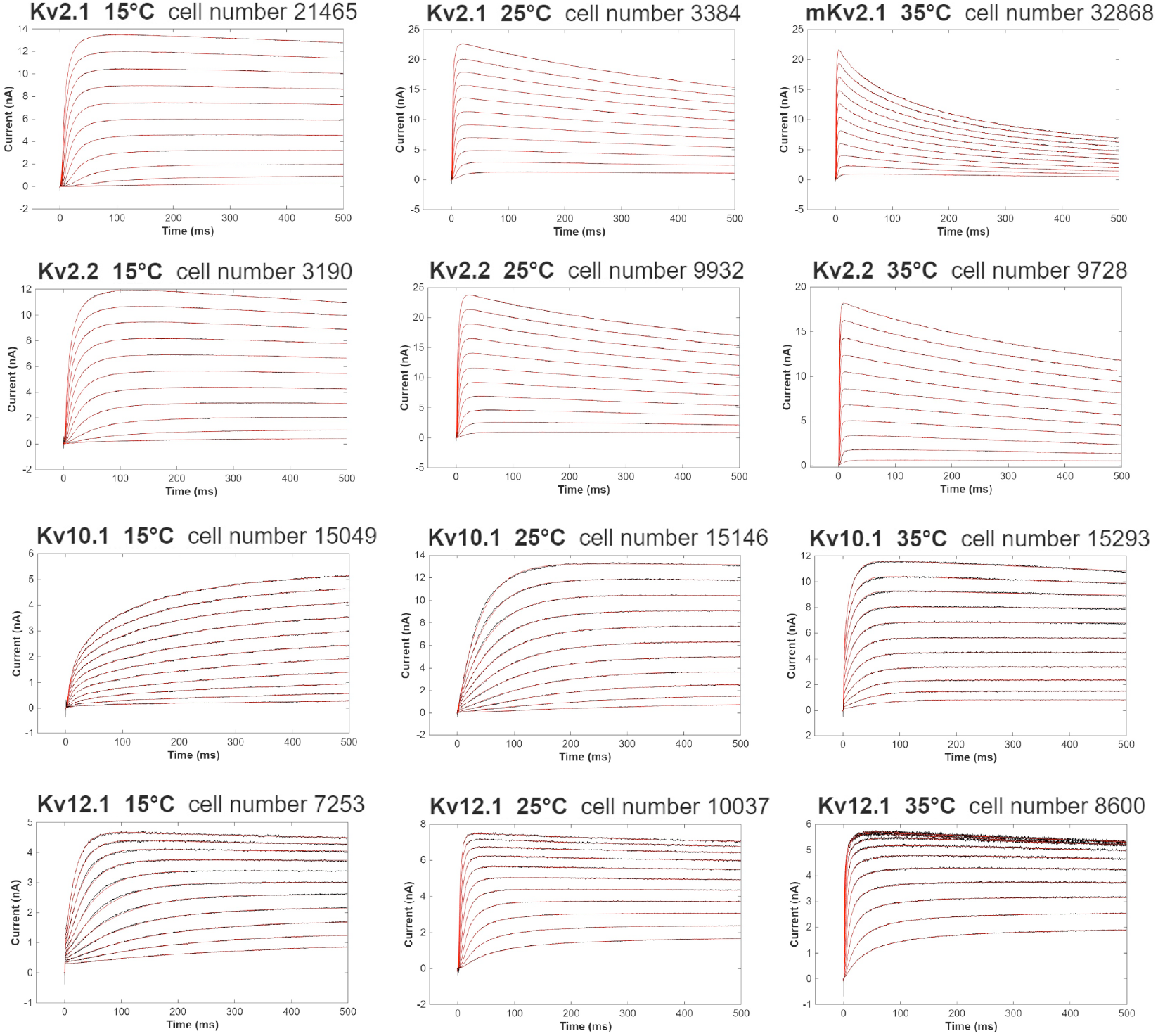
Kv2.1, Kv2.2, Kv10.1, and Kv12.1 activation currents (black curves) fitted by the MS model (red curves).

In most cases, the four-state MS model was sufficient to fit the currents well. When more complex current-curvatures occurred indicating additional states emerged, the five-state scheme was used, fitting Kv1.3 (25 °C and 35 °C), Kv1.4 (35 °C), Kv1.6 (15 °C), Kv1.7 (25 °C), Kv3.2 (35 °C), Kv3.3 (15 °C, 25 °C, and 35 °C), Kv3.4 (15 °C, 25 °C, and 35 °C), and Kv4.3 (35 °C) channel currents. Note that these currents, except Kv1.3, Kv3.2, Kv3.3, and Kv3.4 currents, may also be fitted approximately by the four-state scheme but at low-level accuracies. These indicate that the four-state model-scheme can be used to compare most Kv channel activation-currents examined in this study. If we were to compare more Kv channel currents, the time period can be reduced so that fewer states can emerge that may follow the same kinetic scheme. We also notice that some channel currents are not fitted well even using the five-state scheme (e.g., Kv3.4 currents), yet more kinetic-states are needed there.

Currents below 0 mV were usually small, leading to large variations in the fitted parameters, which sometimes showed anomalous behaviors deviating from the regular dependence on voltage, such as those observed in *i*_1_, *i*_2_, *i*_3_, and *i*_4_ parameter plots (Figs. S1-S18). Here, the fitting-qualities are checked by the RMSE values (Root Mean Squared Error, provided as one of the goodness-of-fit parameters in Matlab software), which are plotted in Fig. 6. Generally, curve fitting at low-voltages can incur large errors (large RMSE values), like those occurred at -20 mV and -10 mV (Fig. 6A and 6B). These errors can result from large noises associated with small-scale currents that increase the uncertainties in curve fitting. Such errors can decrease (gradually) and approach constant levels at high-voltages (Fig. 6C-6E).

**Figure 6.**
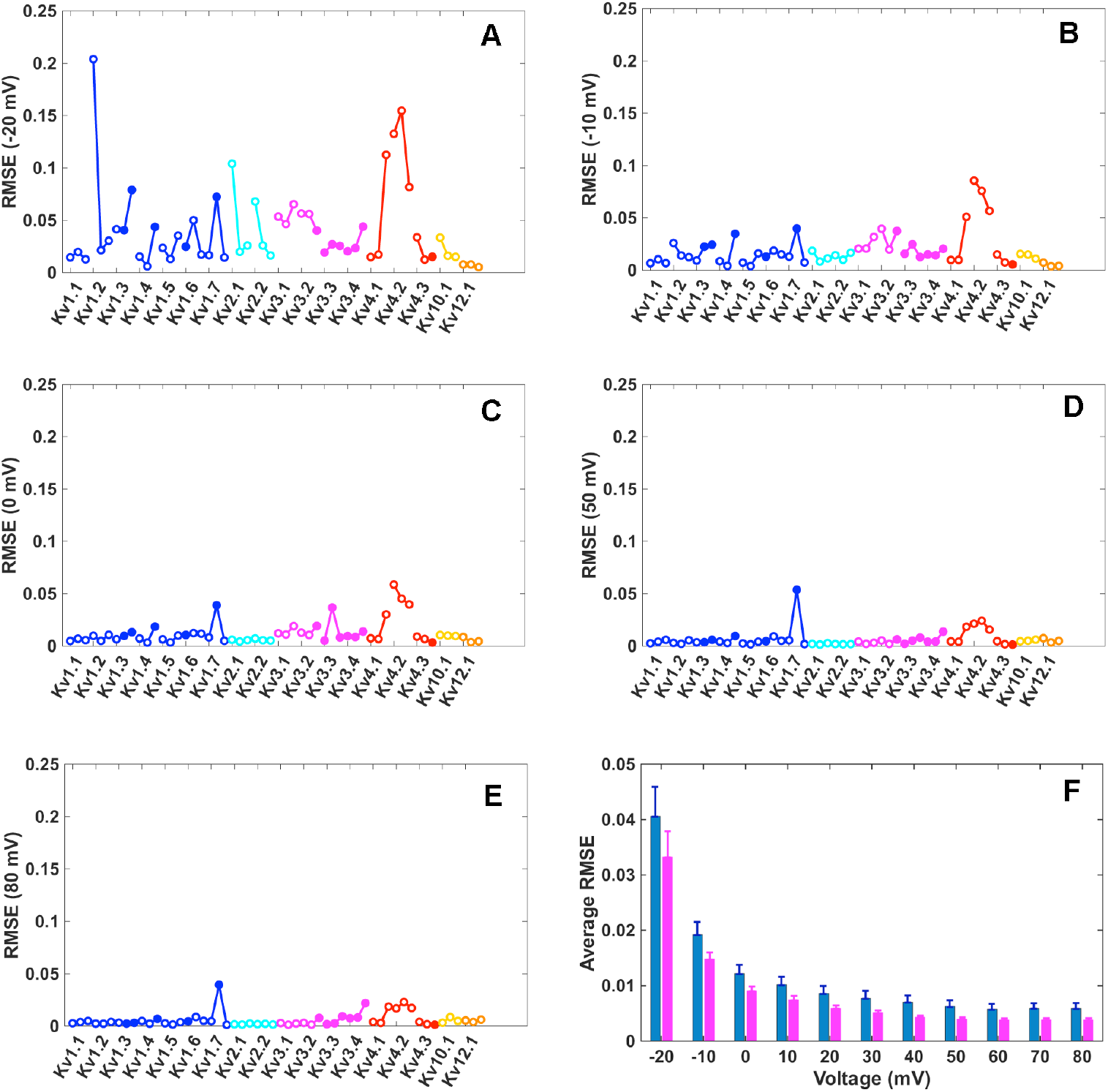
Comparisons of RMSE values obtained from fitting Kv channel activation-currents at different voltages. RMSE values of fitting 54 cell-currents (18 Kv channel currents at three temperatures) recorded at -20 mV (A), -10 mV (B), 0 mV (C), 50 mV (D), and 80 mV (E) are shown individually. In these subplots, each Kv subtype labeled under the x-axis is associated with three RMSE values, represented by three circles connected by lines, corresponding to results obtained at three temperatures 15 °C, 25 °C, and 35 °C, respectively. Open and filled circles represent results from the four-state and five-state scheme fits, respectively. Colors are used to distinguish Kv channel subfamilies, blue for Kv1, cyan for Kv2, magenta for Kv3, red for Kv4, yellow for Kv10, and orange for Kv12 channels, respectively. (F) Average RMSE values calculated for Group A (all Kv channels examined, blue color) and Group B (see text for Group B definition, magenta color) channels. Error bars denote standard errors.

At 80 mV, only few results show large RMSE values (Fig. 6E). Those associated with Kv1.7 (25°C) and Kv3.4 (35 °C) results indicate imperfect fitting. Indeed, these channels require additional states in modeling, where even the five-state scheme is insufficient to fit their currents. Kv4.2 (15 °C, 25 °C, and 35 °C) and Kv4.1 (35 °C) results also demonstrate large RMSE values at 80 mV. These are due to the large noises in the recorded currents (especially for Kv4.2 channels that have small current scales). Here, the RMSE values of all fittings are defined as group A, and those excluding Kv1.7 (25 °C), Kv3.4 (35 °C), Kv4.1 (35 °C), and Kv4.2 (15 °C, 25 °C, and 35 °C) results are defined as group B, and their average values at different voltages are compared in Fig. 6F. Clearly, group B results are more reliable. In addition, the average RMSE values are obviously large at -20 mV and -10 mV. For example, averaging all group-B RMSE values over all voltages leads to the value 0.0087, and it reduces to 0.0052 when *n*8 and *n*9 data (recorded at -20 mV and -10 mV) are excluded. Besides the large noise at low-voltages, special features in currents may also affect the fitting quality. For example, Kv12.1 (15 °C) cell-recording shows anomalously large currents within 1 ms, making *p*_0_, *q*_0_, and *k*_1_ predictions unreliable.

### Features of MS model parameters

Parameters of the four-state MS model used to fit Kv channels are plotted in Fig. 7, and the relations between *k*_1_ and *p*_0_, *q*_0_, *k*_2_, and *k*_3_ are examined in Fig. 8. Parameters of all fittings can be found in Figs. S1-S18. The linear relations between *k*_1_-*p*_0_ and *k*_1_-*q*_0_ are remarkable (Fig. 8A and 8B), indicating that the concatenated opening of the voltage-sensor controlled gate (modeled by parameters *p*_0_, *q*_0_) prompts quick switching from a non-conducting to a conducting channel state (modeled by parameter *k*_1_).

**Figure 7.**
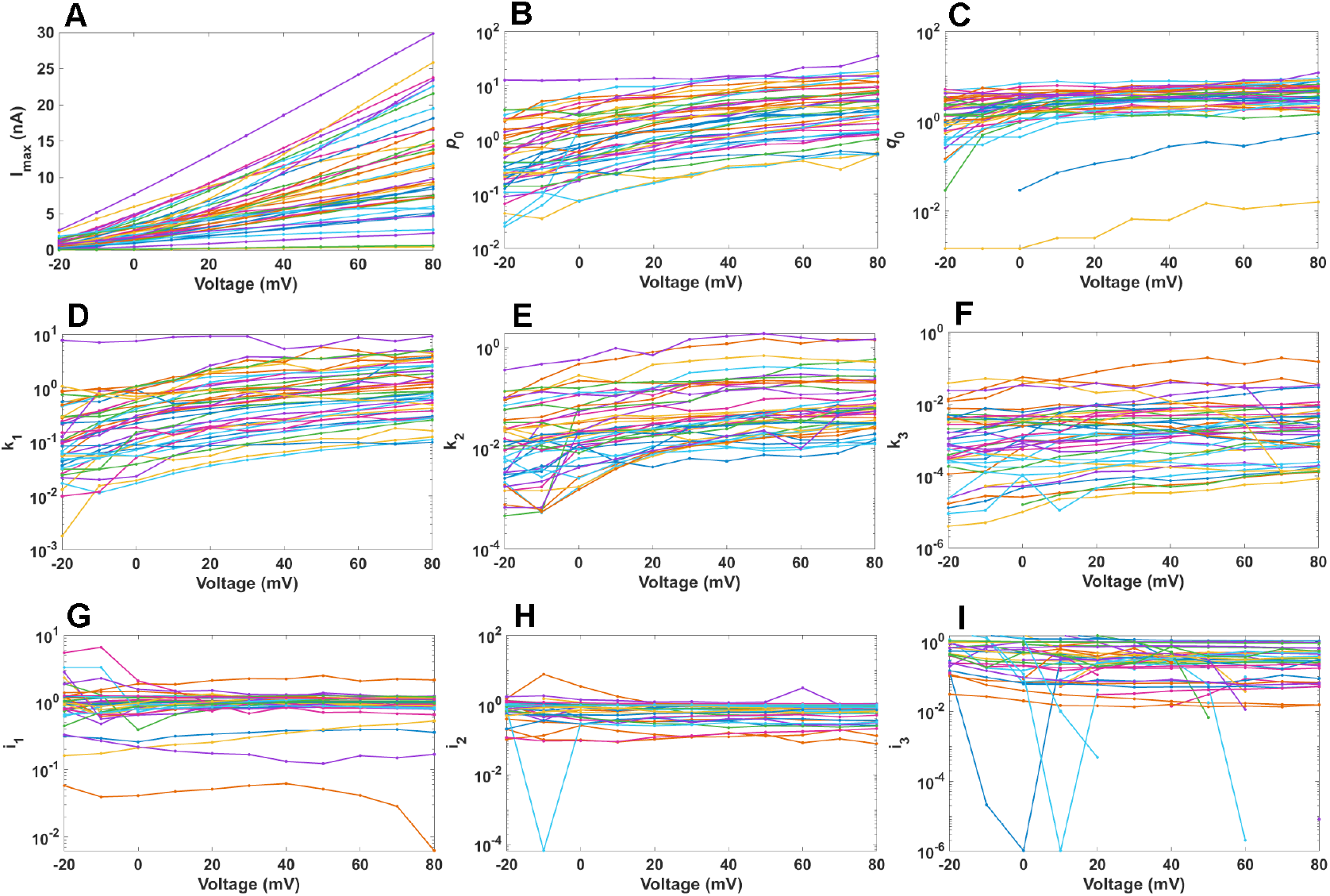
MS model parameters obtained for fitting Kv channel currents using the four-state scheme. The 9 parameters are I_max_ (A), *p*_0_ (B), *q*_0_ (C), *k*_1_ (D), *k*_2_ (E), *k*_3_ (F), *i*_1_ (G), *i*_2_ (H), and *i*_3_ (I).

**Figure 8.**
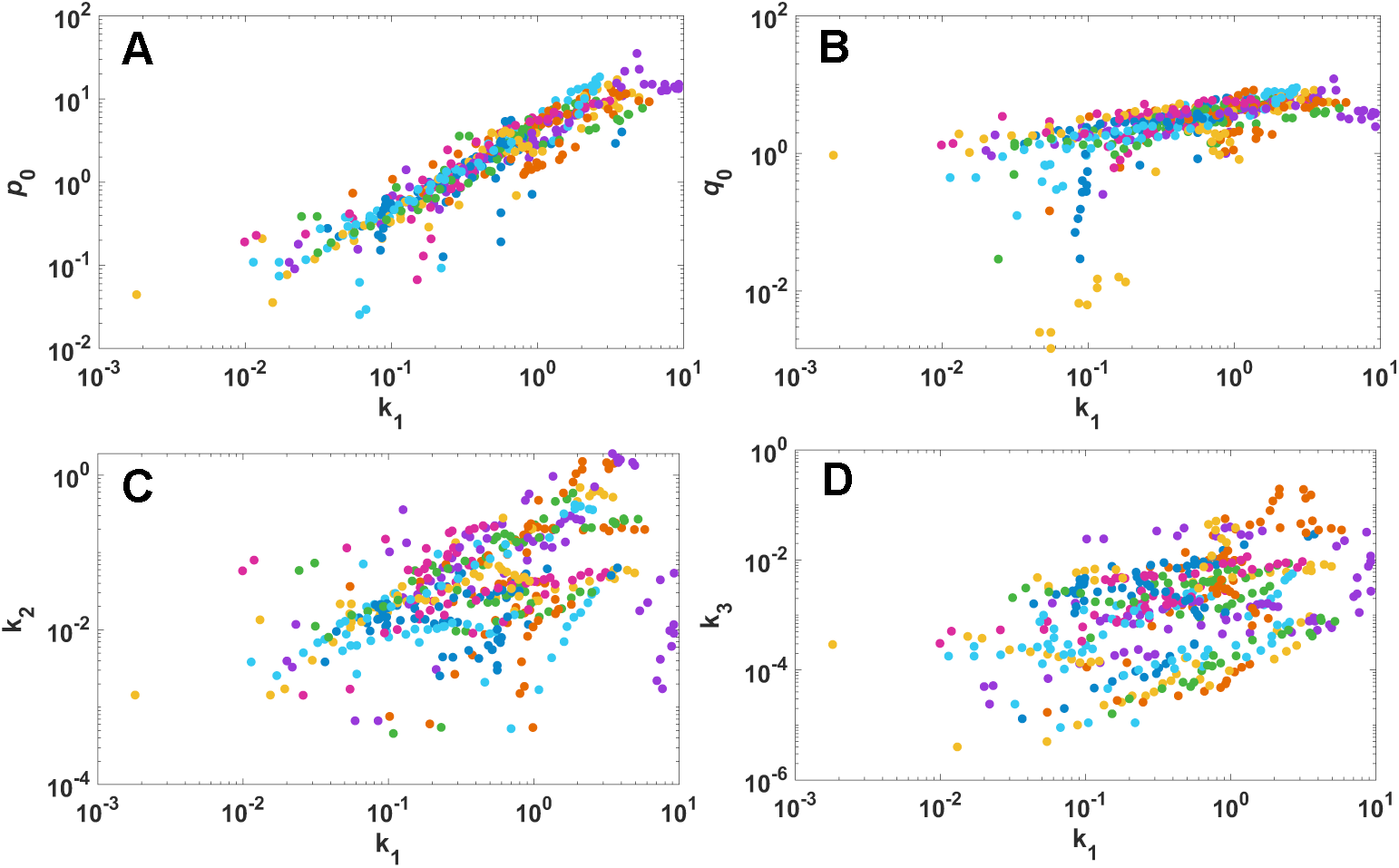
Relations of MS model parameters in the four-state kinetic scheme. Relations of *k*_1_-*p*_0_, *k*_1_-*q*_0_, *k*_1_-*k*_2_, and *k*_1_-*k*_3_ are plotted in (A), (B), (C), and (D), respectively.

MS model parameters demonstrate simple voltage dependence (Figs. 7 and S1-S18). The linear dependences of *p*_0_, *q*_0_, *k*_1_, *k*_2_, *k*_3_, and *k*_4_ on voltage were observed in the initial fit of the current traces. Based on these linear dependences, *p*_0_, *q*_0_, *k*_1_, *k*_2_, *k*_3_, and *k*_4_ were further fine-tuned manually by restricting the range of their variations during fitting. In the final results, they usually showed regularly linear dependence on voltage (Figs. S1-S18). Throughout fitting, room was allowed for these parameters to vary around a line (at least ±20%) so that they would disperse around a line if they did not strictly follow a linear dependence on voltage. Whenever this happens, it can indicate an imperfect mechanism described for that channel, but it does not affect the cross-channel comparison via only a pseudo kinetic scheme where the fitting quality is satisfactory.

Parameters *i*_1_, *i*_2_, *i*_3_, and *i*_4_ were free to vary between 0 and 9.9 during the fitting and refining processes unless otherwise stated (see Materials and Methods). This produced large variations on *i_j_* (*j* = 1∼4, see Figs. S1-S18), especially at low voltages where the RMSE values were intrinsically large. We had not manually set *i_j_* values to follow a regular voltage-dependence. Instead, we left them to vary freely so as to obtain a better perception of *i_j_*-voltage relations. We find that *i_j_* values normally follow a linear or exponential dependence on voltage (Figs. S1-S18) despite some irregular values occurred at low voltages. However, we are not certain about the exponential behaviors shown in these plots, because they rely heavily on the low-voltage data, which are both of insufficient amount (no data below -20 mV) and bearing large RMSE values (Fig. 6).

### Kinetic scheme interpreted by model parameters

Parameters obtained through modeling (Fig. 7) can help describe the features of the three states (*B*, *C*, and *D*) employed in this four-state kinetic scheme. The three rate constants are of declining magnitudes, *k*_1_ > *k*_2_ > *k*_3_ (Fig. 7D-7F), showing the chronological occurrence of the three states. Parameter *i_j_* (Fig. 7G-7I) helps describe properties of each state. *i*_1_ values are usually around 1, indicating a fully-open State *B* normally. *i_2_* values are around or below 1, indicating that State *C* can be a secondary open-state with a reduced conductance level or, in some cases, a gradually inactivating state. Note that each state can be a composite of several substates, and its *i_j_* value depends on these substate populations. Yet a set of varying substate populations can produce a gradually changing *i_j_* values over voltages. If the populations of the low-conducting (or even the non-conducting) substates increase, the results can resemble a gradually inactivating state producing a gradually inactivating current. Such explanations apply well to State *D*, where *i*_3_ values are normally small and close to 0, which is a good indication of an inactivating or a gradually inactivating State *D*. However, in some cases (e.g., Kv10.1 channel currents at 15 °C), *i*_3_ ≈ 1, indicating an open State *D*. Thus, state classifications can vary depending on individual *i_j_* values.

It is worth mentioning that, unlike rate constants *k_j_*, parameter *i_j_* values do not follow uniformly decreasing order over three states. Instead, their values depend on individual current behaviors. Three examples are shown in Fig. 9. Here, Kv10.1 channel currents recorded at 15 °C show non-inactivating behavior over 500 ms. This feature was modeled by a set of increasing *i_j_* values (*i*_1_ < *i*_2_ < *i*_3_) over three states (Fig. 9A), producing gradually increasing macroscopic currents. In the second example, Kv2.2 channel currents recorded at 35 °C start to develop inactivation at high voltages, and this was modeled by relatively constant *i*_1_ and *i*_2_ values (remaining at 1) followed by a decreasing *i*_3_ value (Fig. 9B). In contrast, Kv1.4 channel currents recorded at 25 °C show rapid inactivation over a short time. This was modeled by a set of decreasing *i_j_* values (*i*_1_ > *i*_2_ > *i*_3_) over three states (Fig. 9C), producing fast inactivating currents. Thus, the reduced current levels can help deduce channel mechanisms.

**Figure 9.**
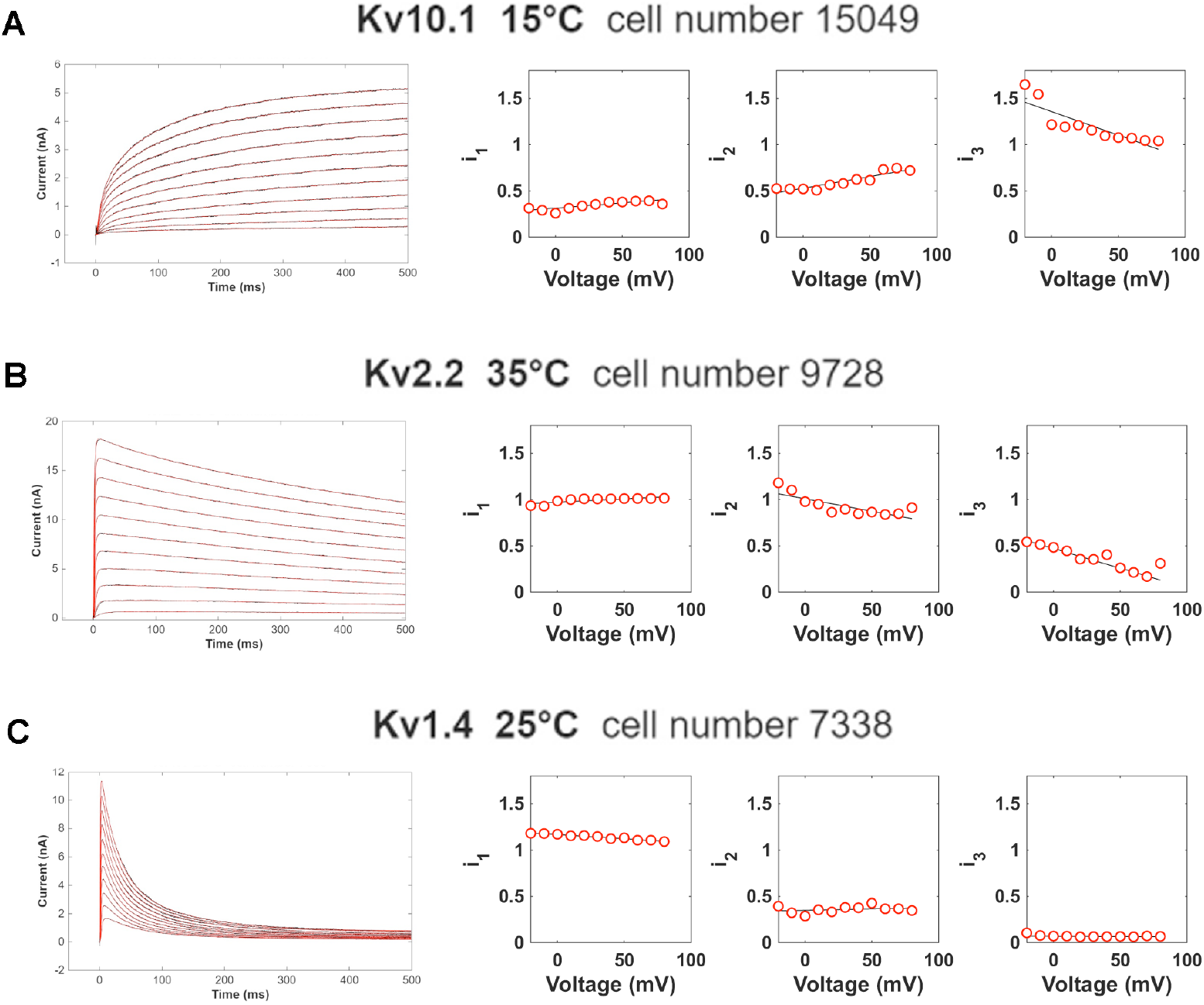
Representative features of the reduced current levels for three typical Kv channel activation currents. The three channel currents, Kv10.1 at 15 °C (A), Kv2.2 at 35 °C (B), and Kv1.4 at 25 °C (C), were all fitted by the four-state kinetic scheme. The fitted current-curves are shown on the left and the model parameters *i*_1_, *i*_2_, and *i*_3_ are compared on the right. Red circles represent parameter values, which are further fitted by linear functions (black lines).

### Temperature dependence of model parameters

Parameters used to model Kv1.1 channel currents at three temperatures (15 °C, 25 °C, and 35 °C) are compared in Fig. 10. Clearly, *p*_0_, *q*_0_, *k*_1_, and *k*_3_ all increase as temperature increases. Besides these, Kv1.1 channel currents (first-row results in Fig. 1) clearly show the development of inactivation as temperature increases. This feature is captured by the change of *i_j_* values over 25 °C to 35 °C, that *i*_2_ and *i*_3_ both decrease while *i*_1_ remains roughly at 1. This means that States *C* and *D* are gradually dominated by the increased populations of inactivating substates, leading to higher levels of inactivation as temperature increases.

**Figure 10.**
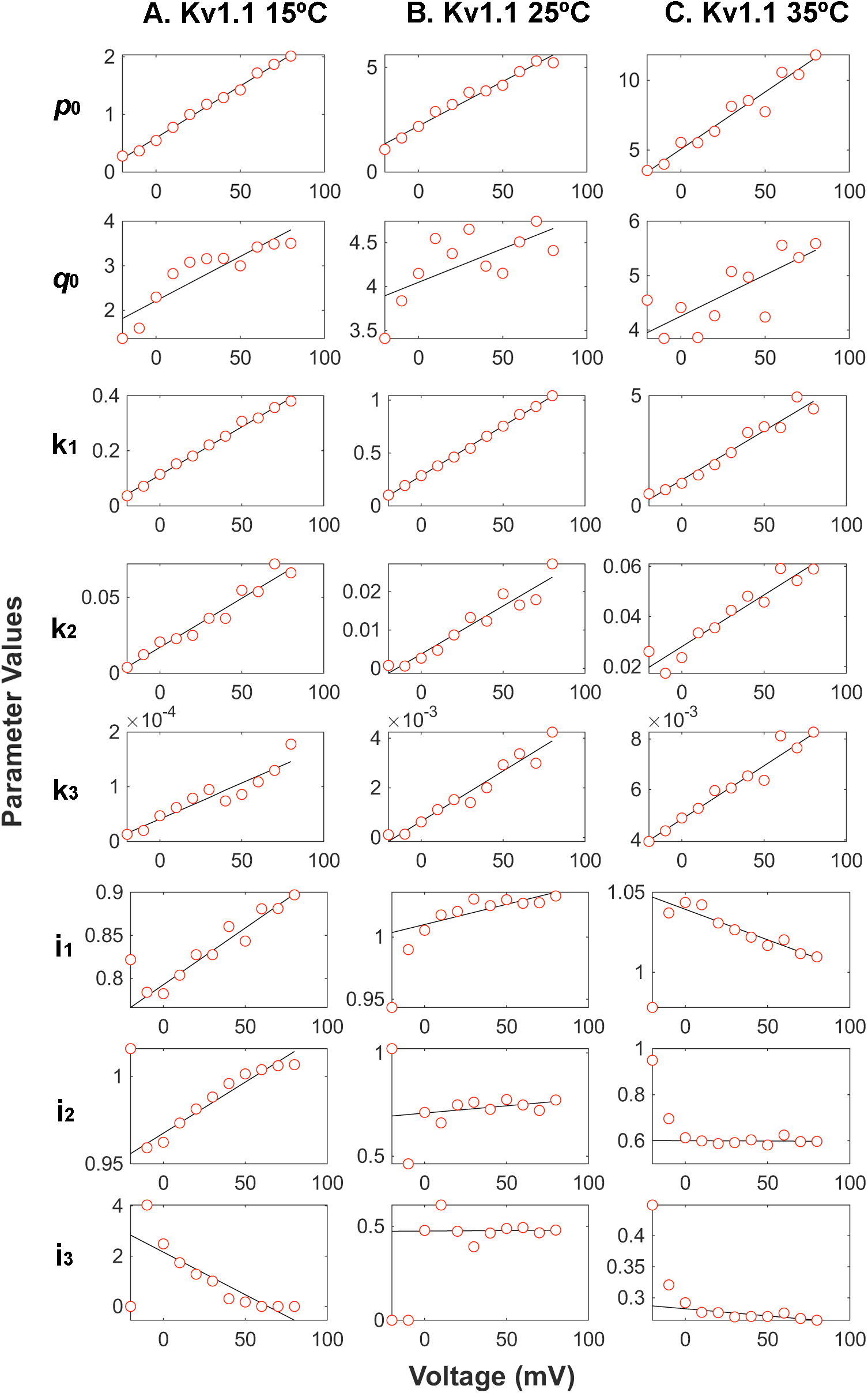
Temperature dependence of the parameters for modeling Kv1.1 channel currents using the four-state scheme. Results at three temperatures 15 °C (A), 25 °C (B), and 35 °C (C) are plotted in each column, where the red circles represent the parameters obtained from the model fitting, and they are further fitted by linear functions (black lines).

Examining all parameters that modeled Kv channel activation currents at 50 mV (Fig. 11), we find that in the majority of channels, *p*_0_, *q*_0_, *k*_1_, and *k*_3_ values increase as temperature increases. These changes can lead to fast opening as well as fast inactivating channel behaviors, which indeed are observed in their currents (Figs. 1-5). Thus temperature can greatly expedite channel opening, activation, and inactivation behaviors, as interpreted by these model parameters.

**Figure 11.**
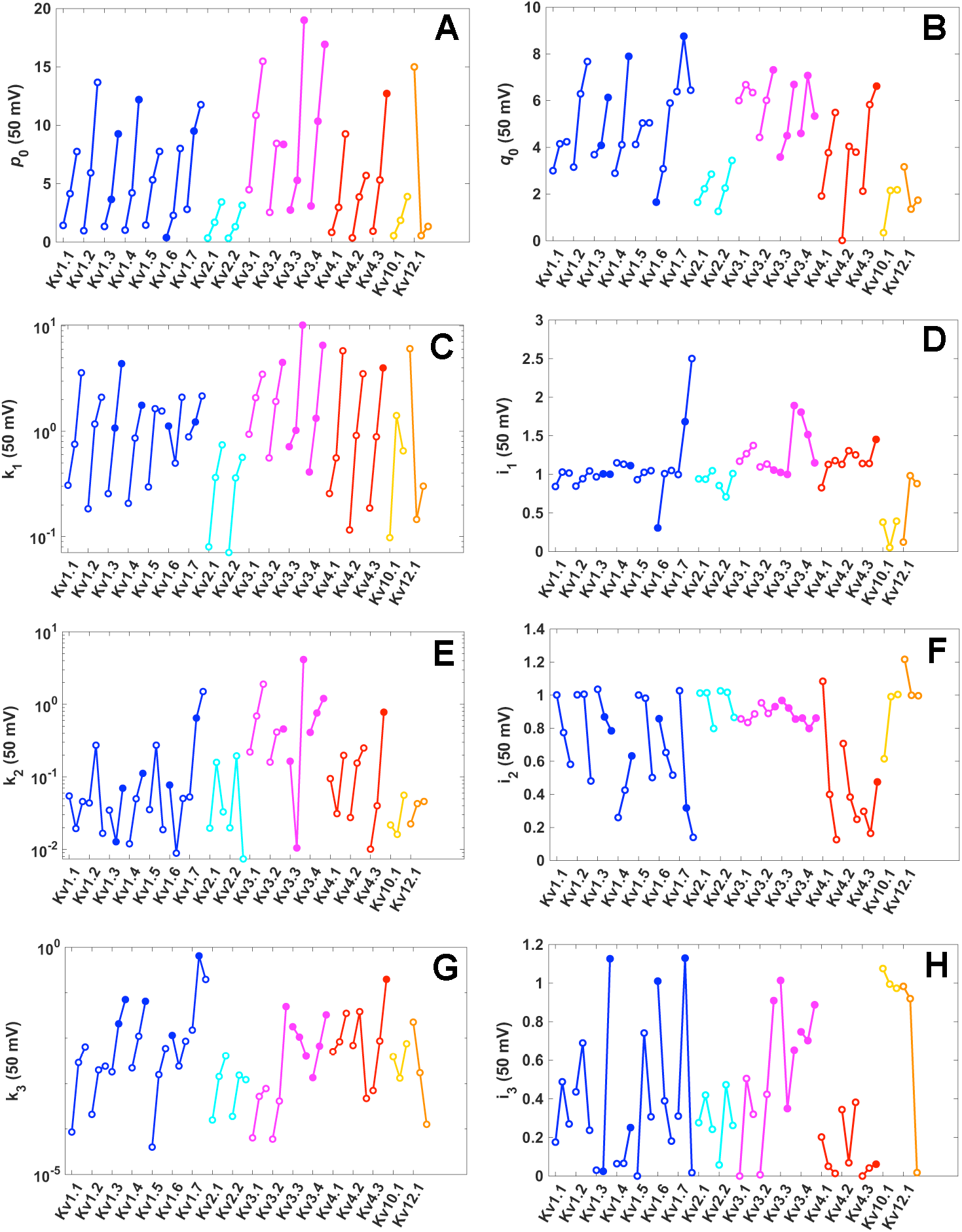
Temperature dependence of MS model parameters for fitting Kv channel activation-currents recorded at 50 mV. Parameters *p*_0_ (A), *q*_0_ (B), *k*_1_ (C), *i*_1_ (D), *k*_2_ (E), *i*_2_ (F), *k*_3_ (G), and *i*_3_ (H) are compared for 18 Kv channels recorded at three temperatures. Each Kv-channel label is associated with three circles connected by lines, representing results at three temperatures 15 °C, 25 °C, and 35 °C, respectively. Open and filled circles represent results from the four-state and five-state scheme fits, respectively. Colors are used to distinguish Kv channel subfamilies, blue for Kv1, cyan for Kv2, magenta for Kv3, red for Kv4, yellow for Kv10, and orange for Kv12 channels, respectively.

Two series of Q_10_ values, Q_10_ (15°C→25°C) and Q_10_ (25°C→35°C), were calculated for parameters *p*_0_, *k*_1_, *k*_2_, and *k*_3_. The results are shown in Fig. 12, where Kv channel subtypes are arranged orderly based on their Q_10_ (15°C→25°C) values. These curves clearly show the different temperature-sensitivities among Kv channel subtypes, that some channels can change their rates of activation and inactivation at low temperatures, whereas others defer these changes at high temperatures. It is also interesting to note that in Fig. 12B and 12C, the two Q_10_ curves show roughly opposite trends of variations, that Q_10_ (25°C→35°C) decreases (approximately) upon the increment of Q_10_ (15°C→25°C). This can indicate a compensating effect in the temperature-induced rate-constant changes, that a large change in one rate constant at one temperature can be accompanied by a small change of the same rate constant at another temperature. Whether this feature applies to other channels requires further studies.

**Figure 12.**
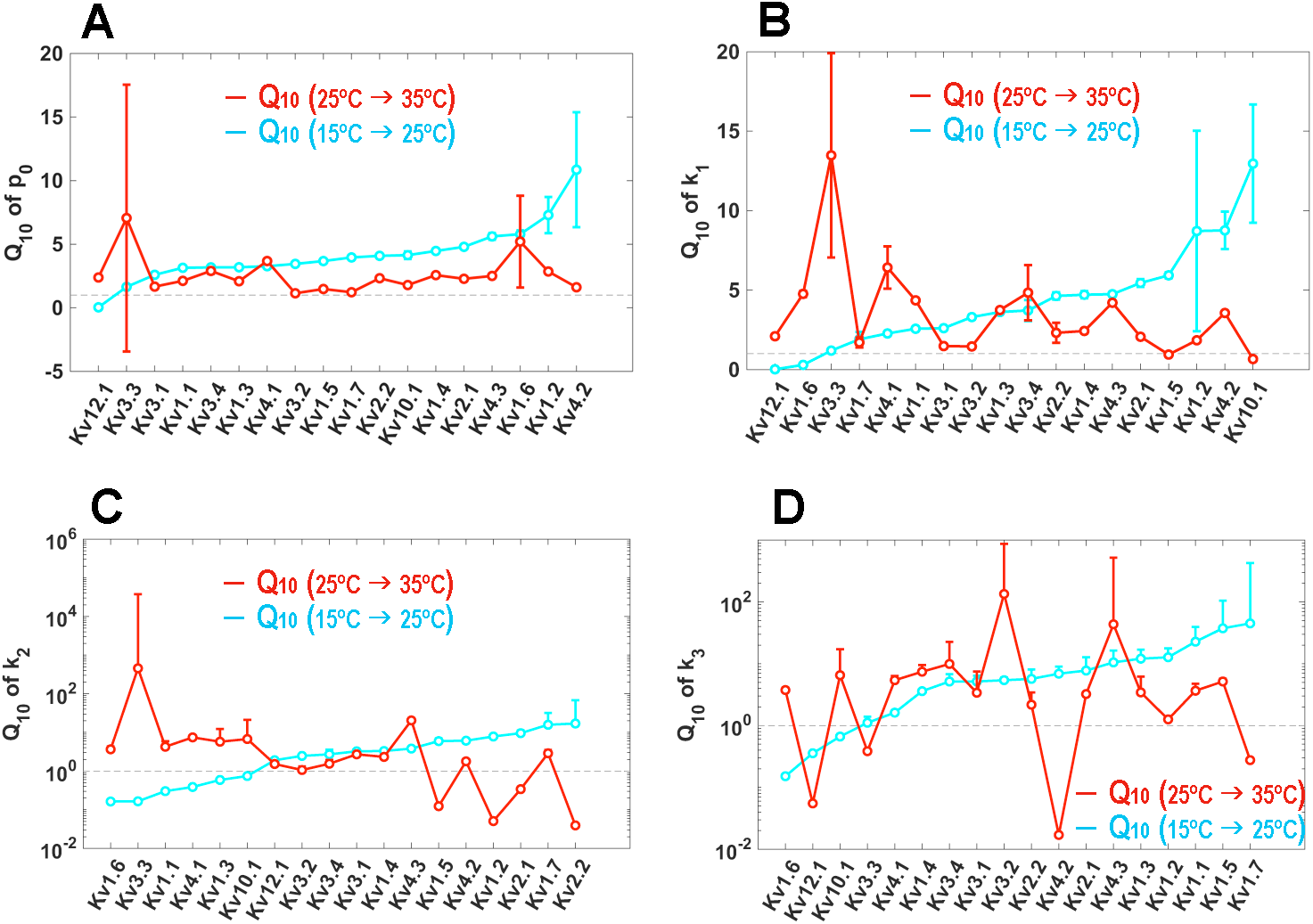
Comparison of Q_10_ values across Kv channels. The Q_10_ values calculated for parameters *p_0_*(A), *k*_1_ (B), *k*_2_ (C), and *k*_3_ (D) are compared for 18 Kv channels. The cyan curves represent Q_10_ (15°C→25°C) and the red curves represent Q_10_ (25°C→35°C). In each subplot, 18 Kv-channel subtypes are arranged orderly based on Q_10_ (15°C→25°C) values calculated for that parameter. Gray dashed lines mark the location of value 1. Each circle represents an averaged Q_10_ value (averaged over nine voltages, from 0 mV to 80 mV). The error bars denote standard errors. In (C) and (D), where the logarithmic scale is used for y-axis, only the upper levels of the error bars are shown.

## Discussion

The gating factor defined by Eq. 4 is indispensable to MS model. It helps fit the beginning period of currents accurately. We compare the effects of modeling Kv channel currents with and without incorporating this gating factor in Fig. 13, where the results in the first and last columns both employed four states whereas those in the middle column employed five states in modeling. It is clear that incorporating the gating factor greatly improves the fitting qualities (last-column results in Fig. 13), which are even better than those of the five-state modeling without the gating factor (middle-column results in Fig. 13).

**Figure 13.**
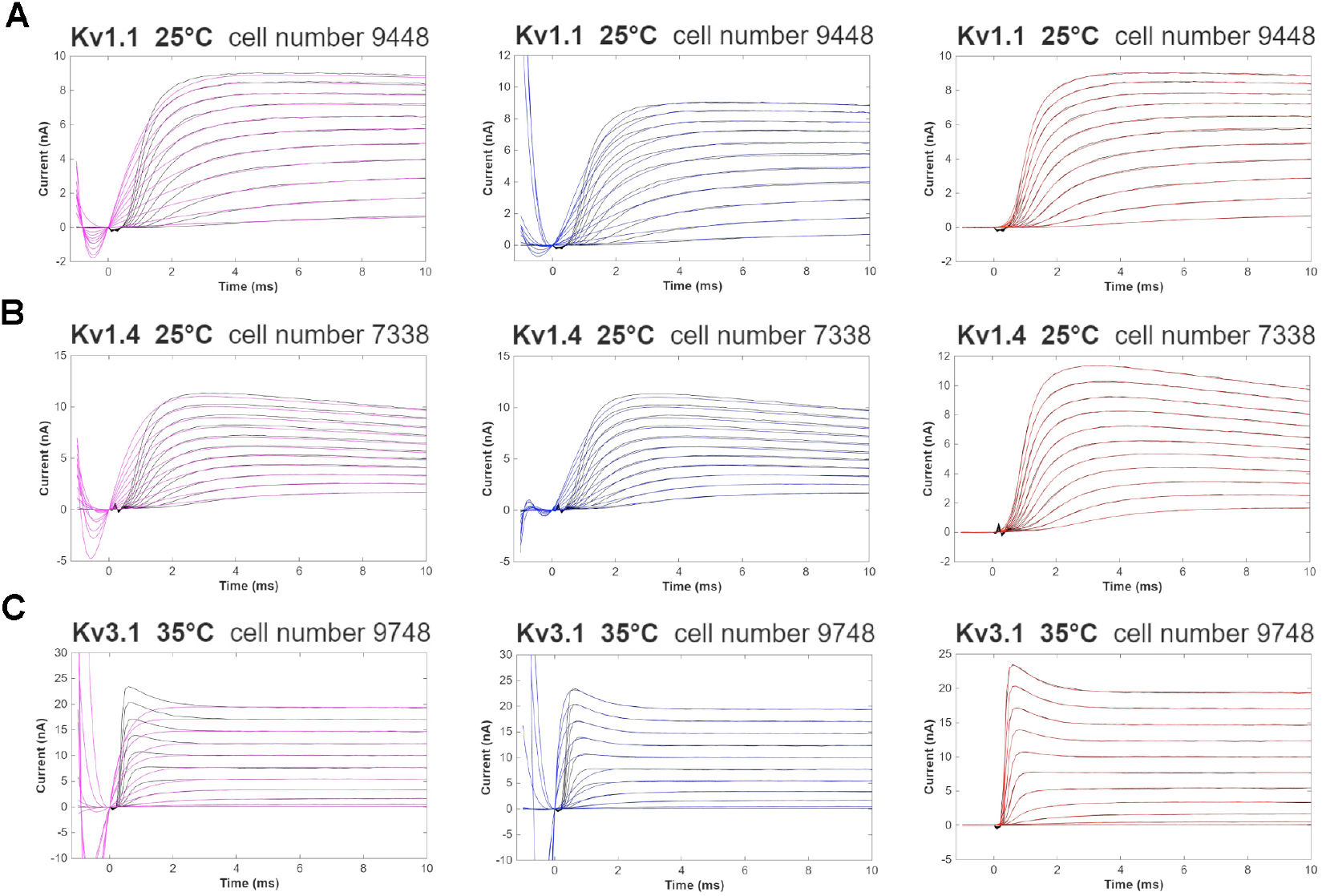
Improved fitting qualities by employing the gating factor. Three channel currents (black curves), Kv1.1 at 25 °C (A), Kv1.4 at 25 °C (B), and Kv3.1 at 35 °C (C), were fitted by the four-state scheme without the gating factor (first-column results, magenta color), the five-state scheme without the gating factor (second-column results, blue color), and the four-state scheme with the gating factor (third-column results, red color). The results at the beginning 10 ms are compared here.

The pseudo four-state kinetic scheme greatly facilitates cross-channel comparisons. Some examples are shown in Fig. 14A-14D, where similar but cross-Kv channel currents (rescaled by their maximum values I_max_) over a range of voltages are identified by comparing their four-state MS model parameters. These identifications are based on the assumption that similar currents are usually modeled by similar *k*_1_, *k*_2_, *k*_3_, *i*_1_, *i*_2_, and *i*_3_ kinetic parameters. Following this rule, similar currents are identified, grouped, and shown in Fig. 14B-14D. For the currents showing little or no inactivation behaviors, such as those in Fig. 14A, their *i*_3_ values can vary greatly. Thus, *i*_3_ value is not considered important in identifying currents shown in Fig. 14A. However, *i*_3_ value must be considered if one wants to compare the tiny differences towards the end of these curves where the inactivation starts to develop.

**Figure 14.**
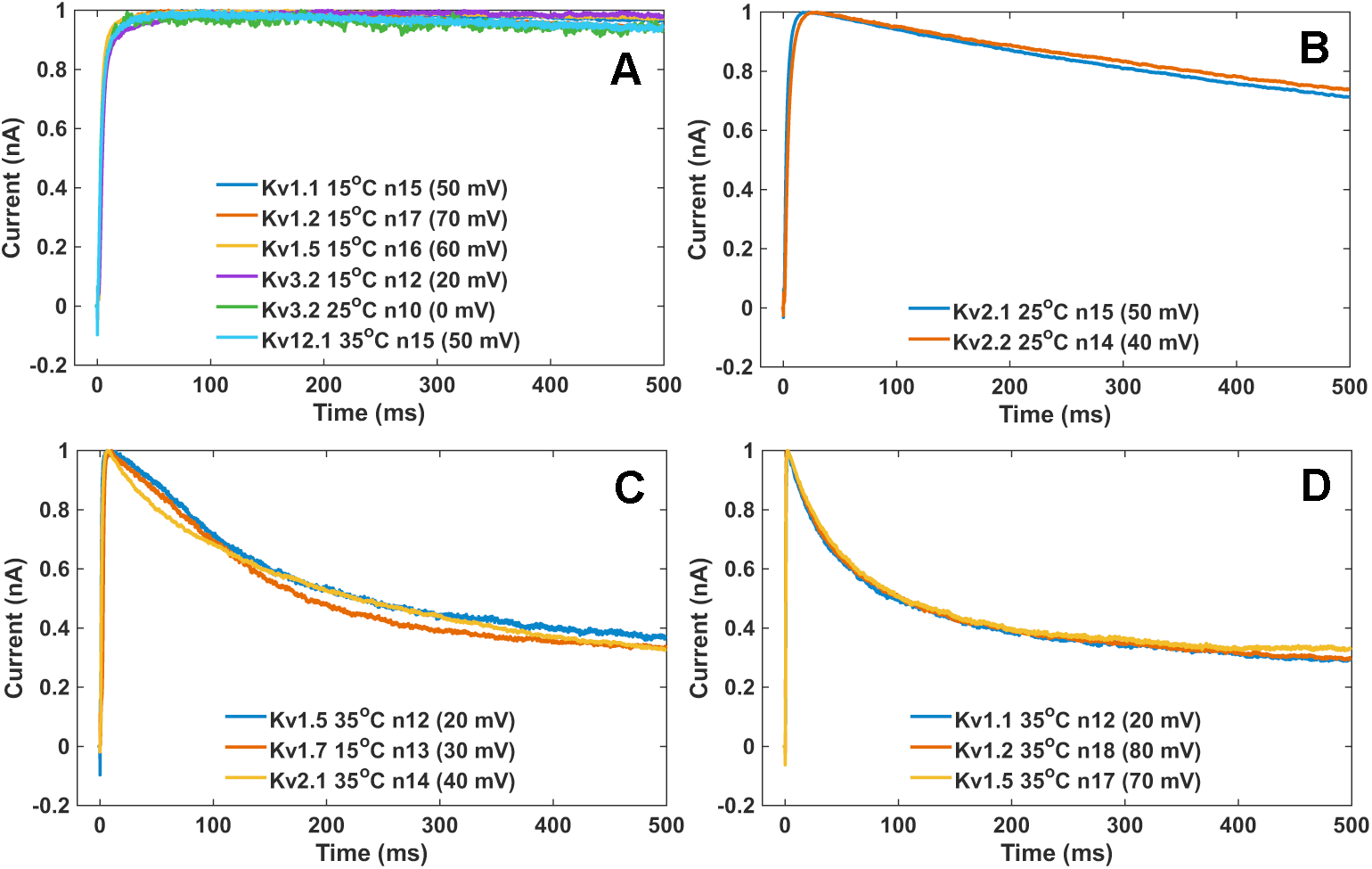
Comparisons of cross-Kv channel activation currents. Similar but cross-Kv channel currents (rescaled by their maximum current values I_max_) are identified by comparing MS model parameters, and they are grouped and shown in (A-D).

MS model parameters help examine a set of regularly varying currents across Kv channels. Three groups of varying currents are shown in Fig. 15, which are all cross-channel currents recorded at 50 mV. They are grouped by channel-activation and inactivation features. Figure 15A shows a set of fast-inactivating currents, which are modeled by a set of obviously increasing *k*_1_, *k*_2_, and *k*_3_ values, relatively constant *i*_1_ values, roughly decreasing *i*_2_ values, and slightly decreasing *i*_3_ values (Fig. 15B). Note that *i*_1_, *i*_2_, and *i*_3_ values are of decreasing magnitudes, i.e., *i*_1_ > *i*_2_ > *i*_3_, producing inactivating currents. The increment of *k_j_* over channels expedites the speed of inactivation. These features are also found in the currents plotted in Fig. 15C, which show similar parameter variations (Fig. 15D) except that *k*_2_ and *k*_3_ become smaller and *i*_2_ becomes larger, resulting in slower speed of inactivation. A set of non-inactivating currents (Fig. 15E) show different features of parameter changes (Fig. 15F). Here, *i*_2_ is larger than *i*_1_. *i*_3_ value remains at 1 for non-inactivating currents and drops below 1 upon the development of inactivation (last case in Fig. 15F, Kv1.1 current at 15 °C). Therefore, the four-state MS model parameters are effective in comparing and characterizing the features of Kv channel activation currents.

**Figure 15.**
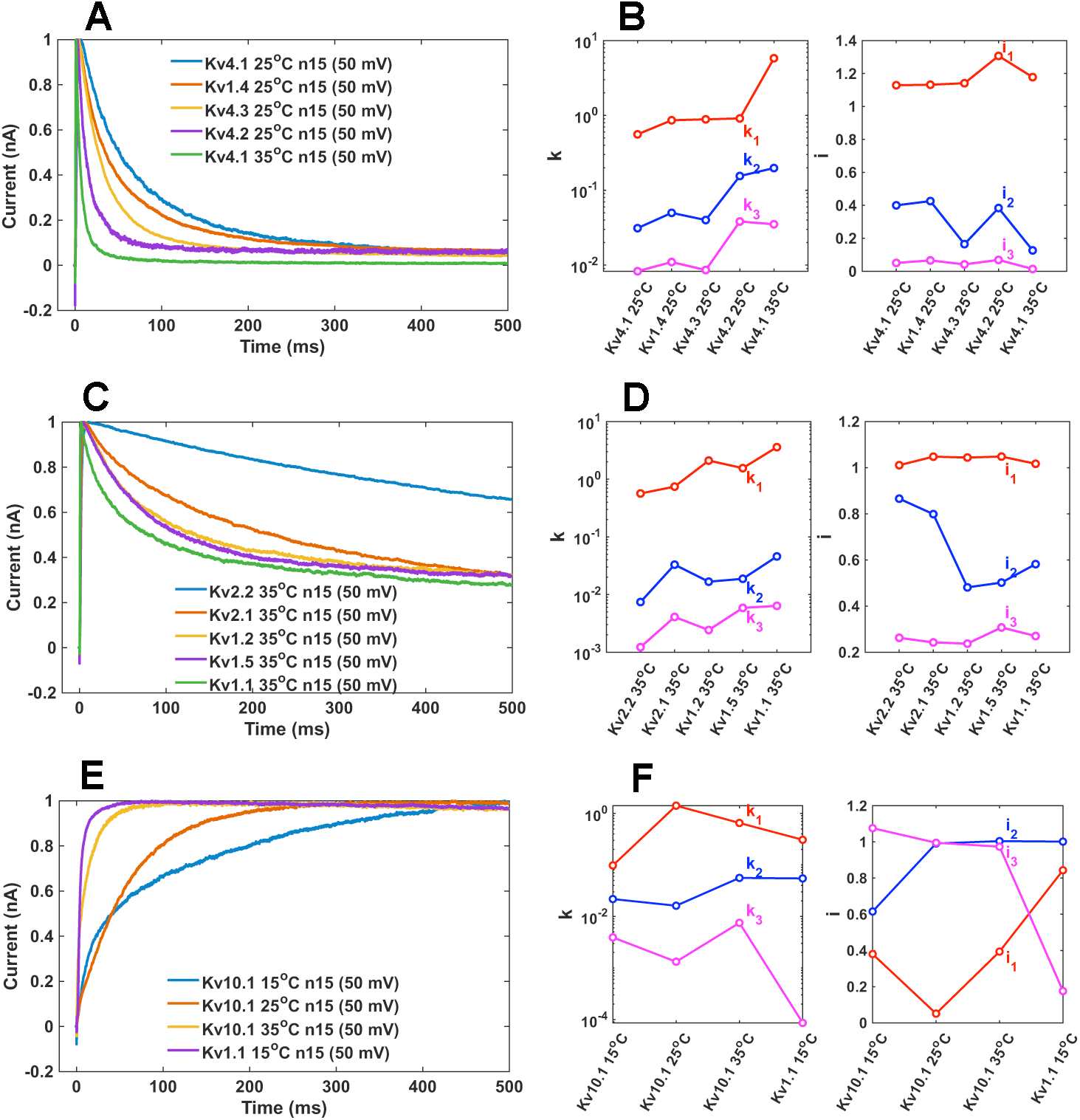
Features of regularly varying Kv channel currents described by MS model parameters. A group of fast inactivating Kv channel currents (A) demonstrate regular changes in their model parameters (B). A group of inactivating Kv channel currents (C) also show regular changes in their model parameters (D). A group of non-inactivating Kv channel currents (E) show different variations in their model parameters (F). Currents were all rescaled by their maximum values I_max_.

One limitation of this work is the identifiability of model parameters, which is a common issue in modeling ion-channel currents^66^. We cannot guarantee that all model parameters obtained here are at their optimal values, and advanced parameter-estimation methods may be investigated in the further.

In conclusion, the MS model proposed here has three major features. (1) A gating factor is incorporated, helping reduce the number of non-conducting states while fitting the beginning period of activation currents accurately. (2) Each state, except the first non-conducting state, is associated with a reduced current level *i_j_*. This helps reduce the number of states needed and improves the fitting quality. (3) A general model framework (defined by Eq. 5) is proposed that facilitates the inclusion of more states and state-transitions necessary for developing channel-specific mechanisms.

## Materials and Methods

Kv channel activation currents recorded at three temperatures (15 °C, 25 °C, and 35 °C) were obtained from Channelpedia website (https://channelpedia.epfl.ch), where 18 Kv channel subtypes were selected, including Kv1.1, Kv1.2, Kv1.3, Kv1.4, Kv1.5, Kv1.6, Kv1.7, Kv2.1, Kv2.2, Kv3.1, Kv3.2, Kv3.3, Kv3.4, Kv4.1, Kv4.2, Kv4.3, Kv10.1, and Kv12.1 channels. The selected current recordings were from rat Kv channels (including a mouse Kv2.1 channel, currents recorded at 35 °C) expressed in CHO cells^62^. Each recording had a unique cell ID number assigned by Channelpedia group, which was also labeled on each result.

The cell-recording data were initially processed by the Matlab and Python codes obtained from Channelpedia website. We used only the second repetition results from the activation-protocol experiments. Each original cell-recording contained 18 activation-current traces, elicited by the stimulation pulses from -90 mV to 80 mV at 10 mV intervals. We had selected only *n*8 to *n*18 current traces of each recording, responding to currents recorded at -20 mV to 80 mV. Currents recorded at low-voltages were not used in order to avoid the ambiguities during fitting. Each stimulation pulse lasted for 500 ms, starting from 100 ms to 600 ms in each recording before returning to the holding potential at -80 mV. In this study, we had selected the time period from 99 ms to 599 ms from the original recordings, corresponding to -1 ms to 499 ms shown in our results. The pre-pulse data (lasted for 1 ms here) of each current curve were averaged, which became a baseline that the whole curve was shifted by this baseline before fitting.

The maximum value over 499 ms time period of each current curve was found and assigned to the model parameter I_max_, and this curve was rescaled by I_max_ prior to fitting. Then, each curve was fitted by Eq. 5 via the Matlab code using the nonlinear least squares method (implemented in the curve-fitting toolbox in Matlab), and the quality of fitting was checked by the RMSE (Root Mean Squared Error) value provided by the Matlab software. In fitting each current trace, the length of data was reduced, that the first 300 data points were always used and the remaining data were picked regularly every 10 points. The negative data points after 0 ms until the occurrence of I_max_ were neglected during fitting. For the rapidly inactivating currents, larger weights were given to the data occurring at earlier time. Therefore, the fitting procedure relied on human intervention.

The five-state MS-model scheme was used for fitting Kv1.3 (25 °C and 35 °C), Kv1.4 (35 °C), Kv1.6 (15 °C), Kv1.7 (25 °C), Kv3.2 (35 °C), Kv3.3 (15 °C, 25 °C, and 35 °C), Kv3.4 (15 °C, 25 °C, and 35 °C), and Kv4.3 (35 °C) channel currents, and the four-state MS-model scheme was used for fitting all other Kv channel currents selected in this study. The results of initial fit always showed that model parameters depended on voltage linearly. Based on these linear behaviors, the parameters were fine-tuned manually in the subsequent rounds of fittings, where parameters *p*_0_, *q*_0_, *k*_1_, *k*_2_, *k*_3_, and *k*_4_ were allowed to change in restricted ranges with boundaries not smaller than ±20% from line behaviors normally. On the other hand, parameters *i*_1_, *i*_2_, *i*_3_, and *i*_4_ were left free to vary in the range of [0, 9.9], even in the manually refining processes, except for the following cases: [0, 19.9] for *i_j_* in Kv1.5 (15 °C), [0, 2.9] for *i_j_* in Kv1.6 (15 °C), [0, 2.9] for *i_j_* in Kv1.7 (35 °C), [0, 1.9] for *i_j_* in Kv2.2 (15 °C), [0, 1.9] for *i_j_* in Kv3.1 (15 °C), [0, 5.9] for *i_j_* in Kv3.4 (15 °C), [0, 1.9] for *i*_4_ (*n*_8_) in Kv1.3 (35 °C), [0, 0.2] for *i*_3_ (*n*_8_) in Kv3.1 (25 °C), and [0, 1.9] for *i_j_* (*n*_8_) in Kv3.2 (15 °C) current models, where *j* = 1∼3 for the four-state scheme and *j* = 1∼4 for the five-state scheme.

Q_10_ value was calculated by the ratio of the rate constant obtained at two temperatures differed by 10 °C. Q_10_ values usually varied over voltages. Here, at a given voltage, two Q_10_ values were calculated: Q_10_ (15°C→25°C) = X (25 °C) / X (15 °C) and Q_10_ (25°C→35°C) = X (35 °C) / X (25 °C), where X was one of the parameters *p*_0_, *k*_1_, *k*_2_, and *k*_3_. Then each set of Q_10_ (15°C→25°C) and Q_10_ (25°C→35°C) values were averaged over nine voltages (from 0 mV to 80 mV) separately, and the averaged Q_10_ values were used for each Kv channel plotted in Fig. 12.

## Supporting Information

**Supplementary Figures S1-S18**. MS model parameters for fitting 18 Kv channel activation-currents recorded at three temperatures are plotted in Figs. S1 to S18 individually, where parameter-voltage relations can be examined.

## Acknowledgements

We gratefully acknowledge the Channelpedia research group for their generously sharing Kv channel experimental data on Channelpedia website (https://channelpedia.epfl.ch). This work was supported by Fudan University.

## Author Contributions

**Conceptualization and modeling**: Di Wu.

**Data preparation**: Shushang Jiang and Yiyun Huang.

**Data analyses**: Di Wu, Shushang Jiang, and Yiyun Huang.

**Writing and reviewing**: Di Wu.

