## Supplementary Figures for "Characterizing features of Kv channel activation currents by a minimal-state kinetic model"

### Supporting Information

**Supplementary Figures S1-S18**. MS model parameters for fitting 18 Kv channel activation-currents recorded at three temperatures are plotted in Figs. S1 to S18 individually, where parameter-voltage relations can be examined.

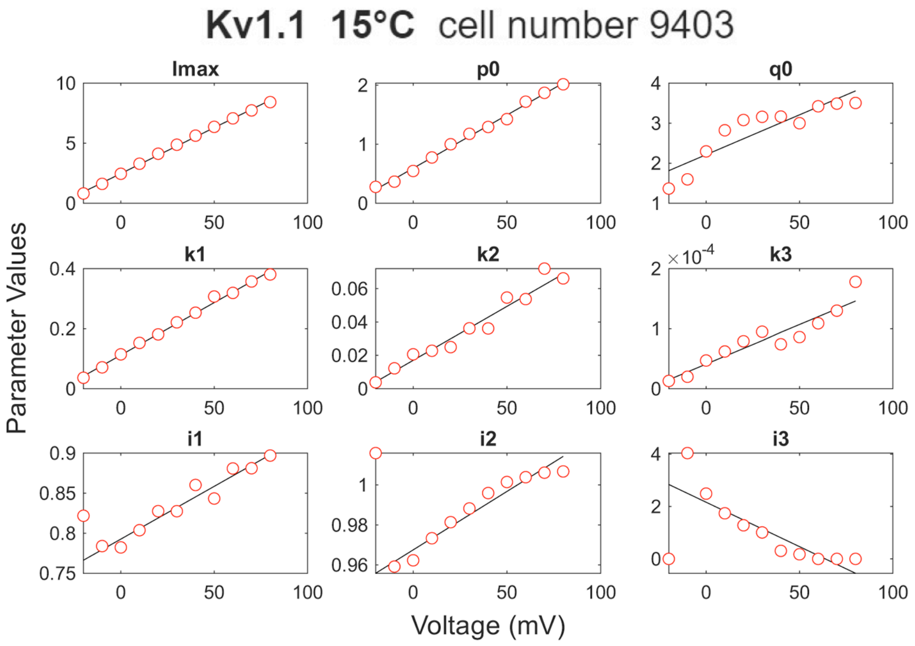


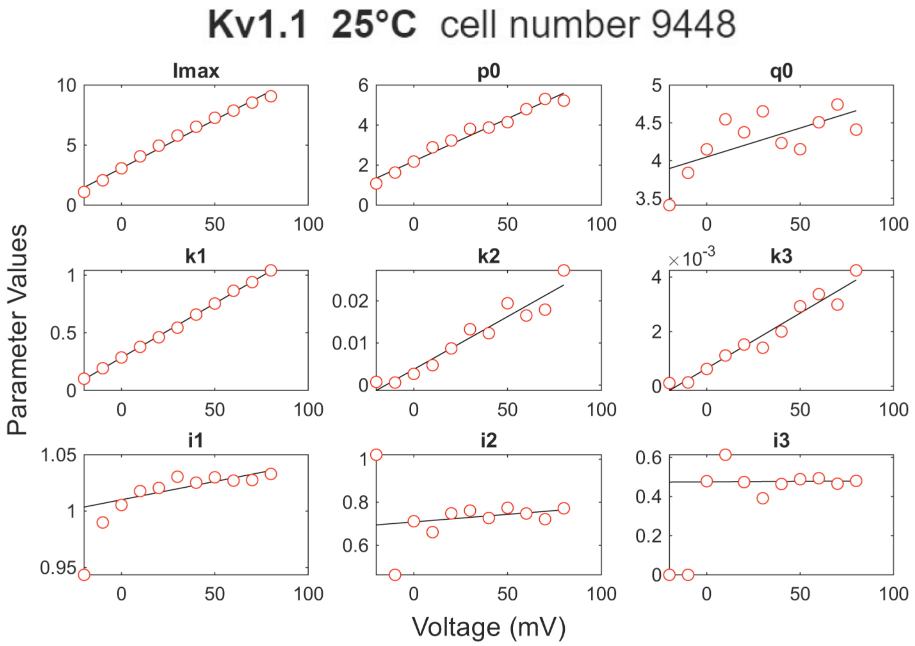


*Figure S1 Continued on next page*

*Figure S1 continued*

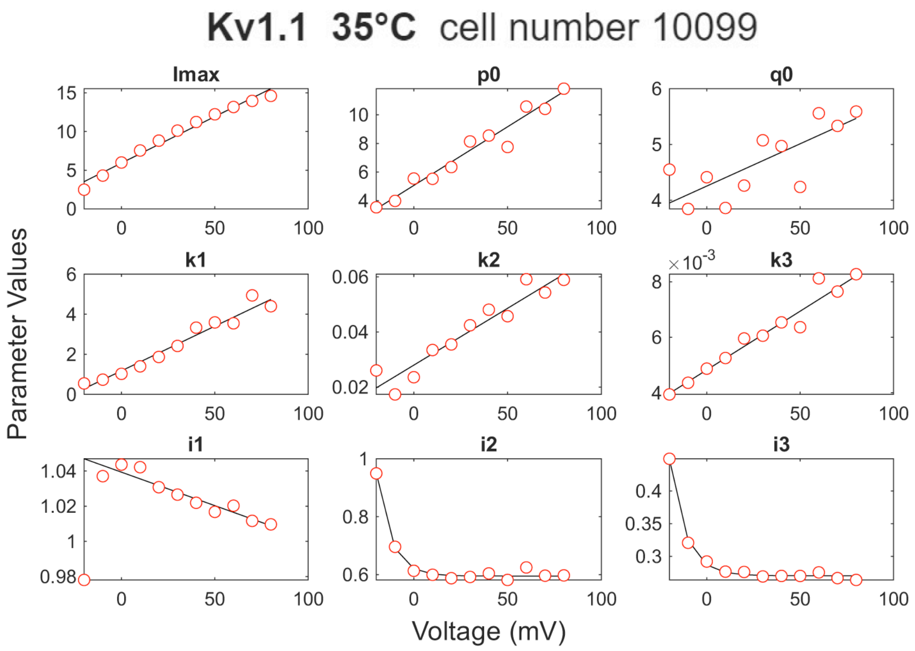


**Figure S1**. MS model parameters for fitting Kv1.1 activation currents at three temperatures. Red circles represent the parameter values obtained from the fitting, which are further fitted by linear functions (black lines) except the Kv1.1 (35 ºC) results where $i_2$ and $i_3$ are fitted by exponential functions.

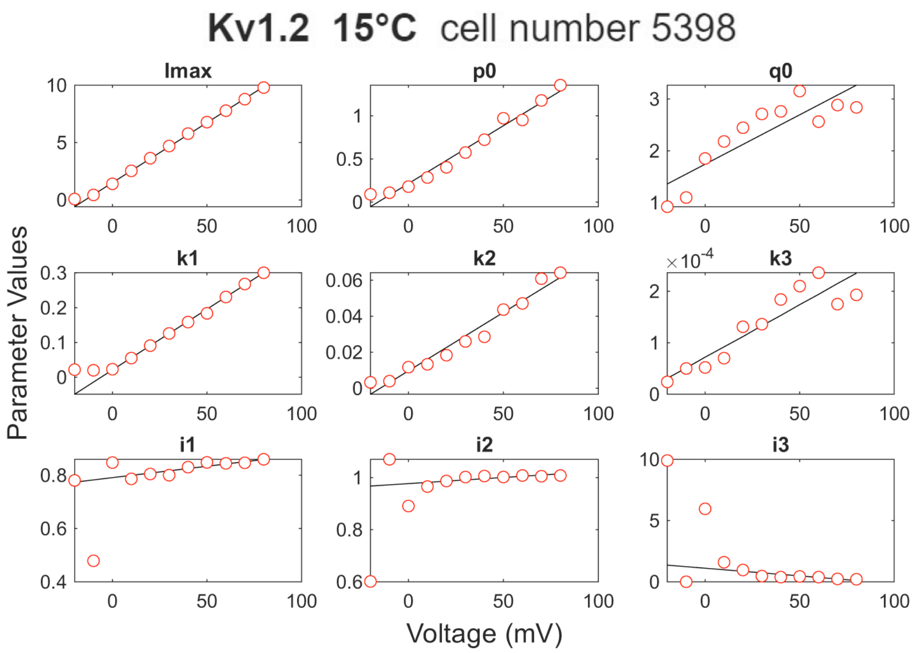


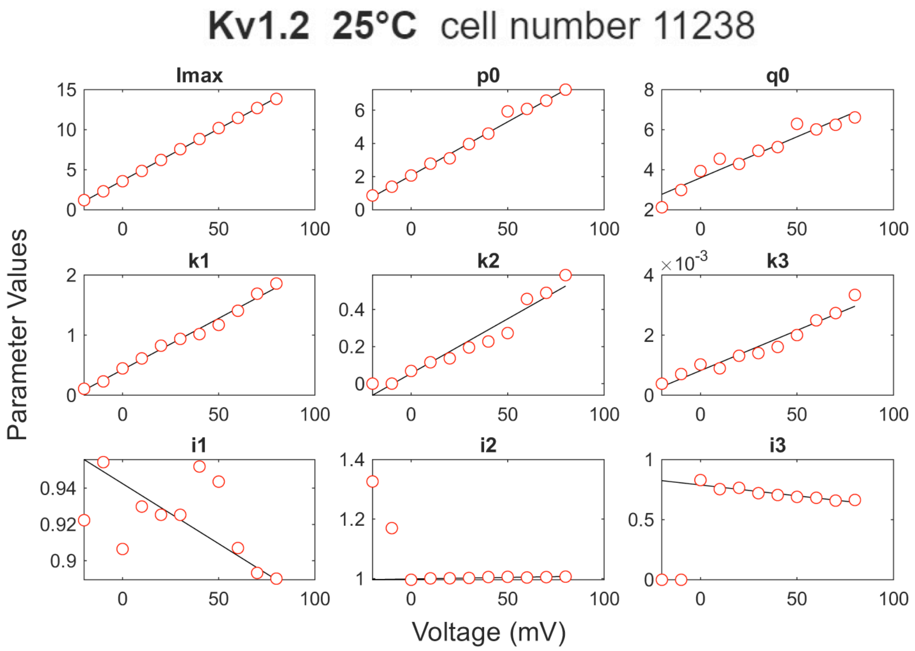


*Figure S2 Continued on next page*

*Figure S2 continued*

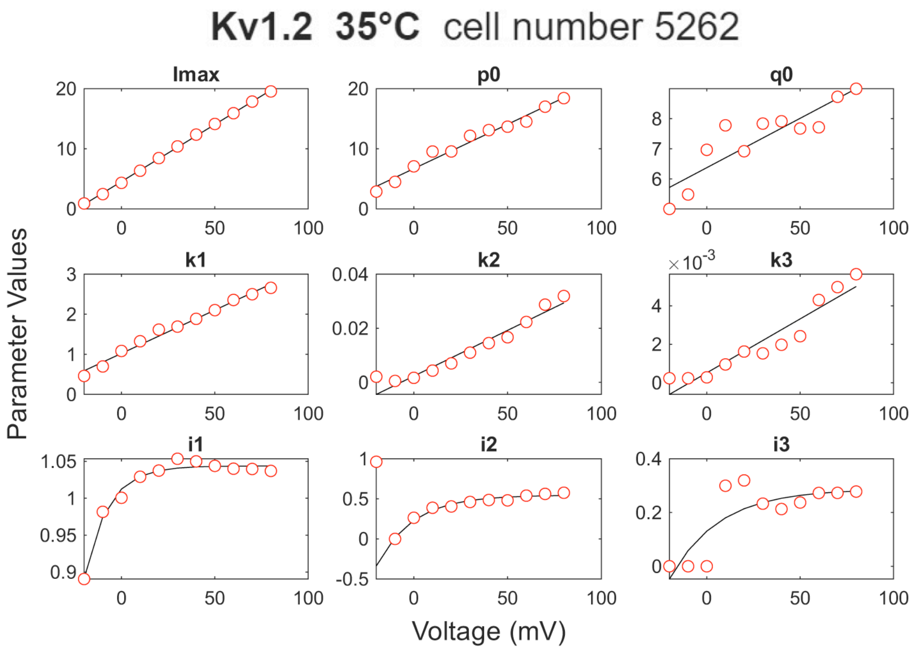


**Figure S2**. MS model parameters for fitting Kv1.2 activation currents at three temperatures. Red circles represent the parameter values obtained from the fitting, which are further fitted by linear functions (black lines) except the Kv1.2 (35 ºC) results where $i_1$, $i_2$, and $i_3$ are fitted by exponential functions.

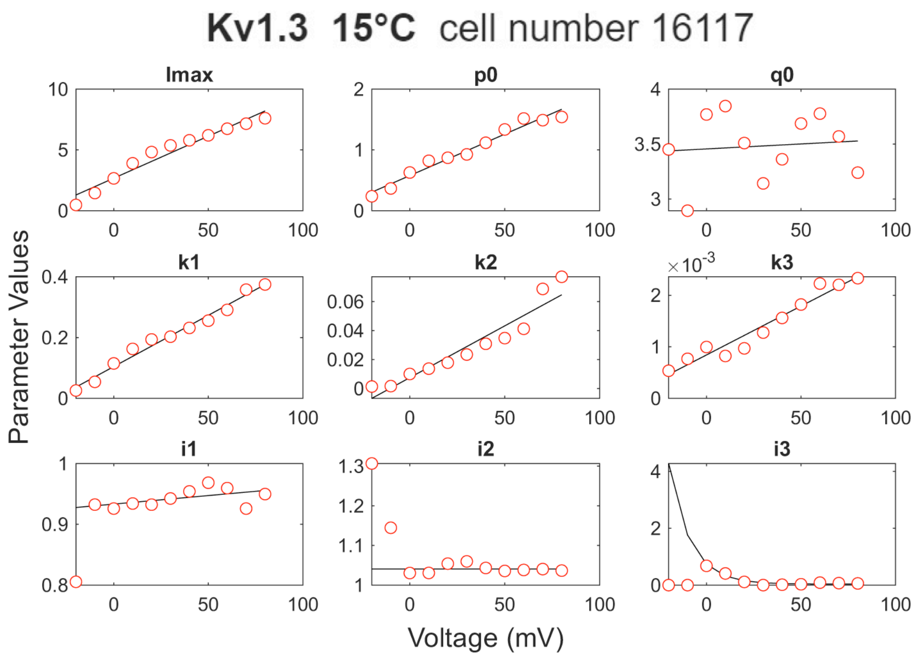


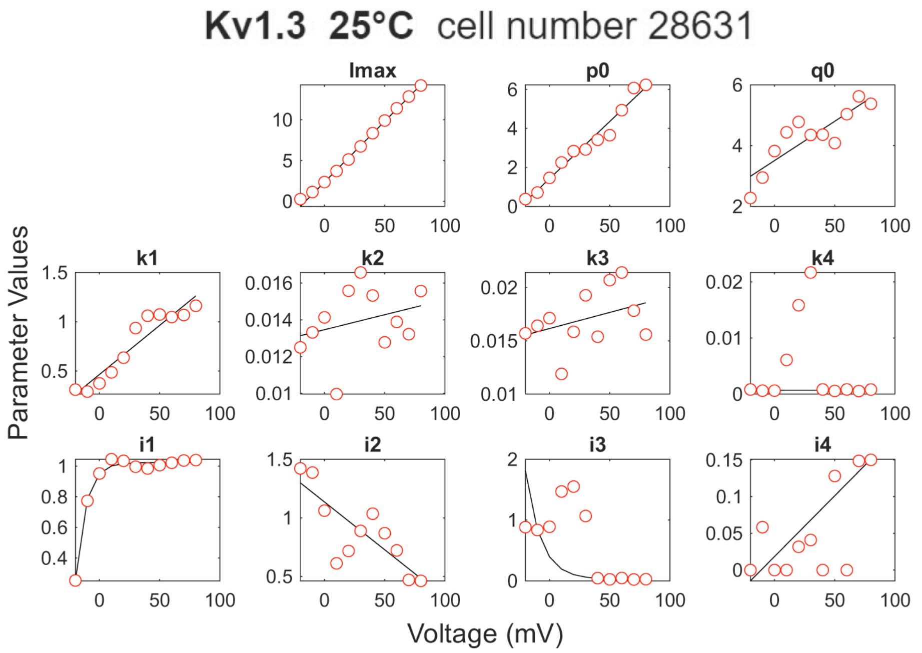


*Figure S3 Continued on next page*

*Figure S3 continued*

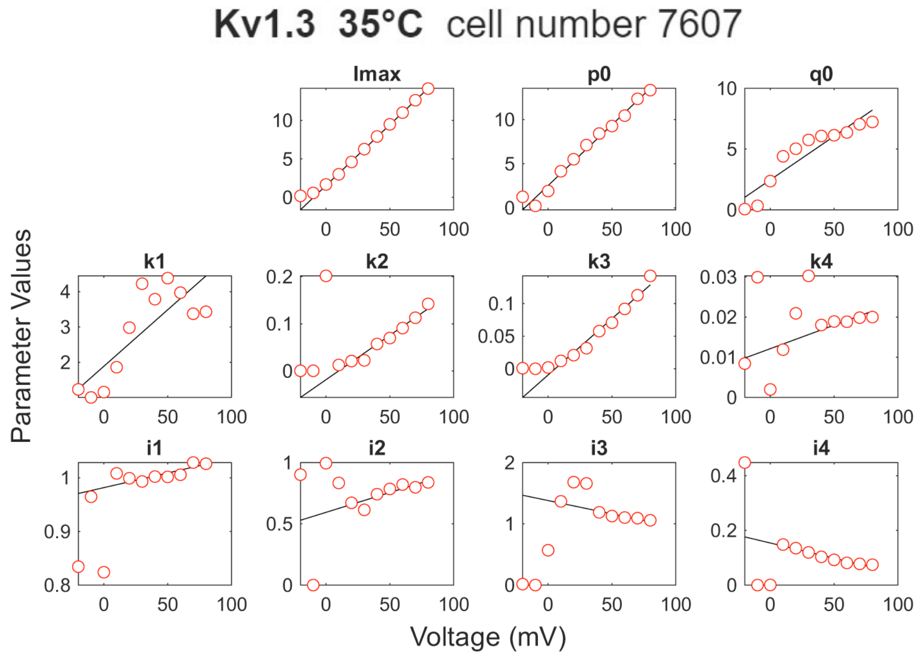


**Figure S3**. MS model parameters for fitting Kv1.3 activation currents at three temperatures. Red circles represent the parameter values obtained from the fitting. They are further fitted by linear functions (black lines) except that $i_3$ of Kv1.3 (15 ºC), and $i_1$ and $i_3$ of Kv1.3 (25 ºC) are fitted by exponential functions.

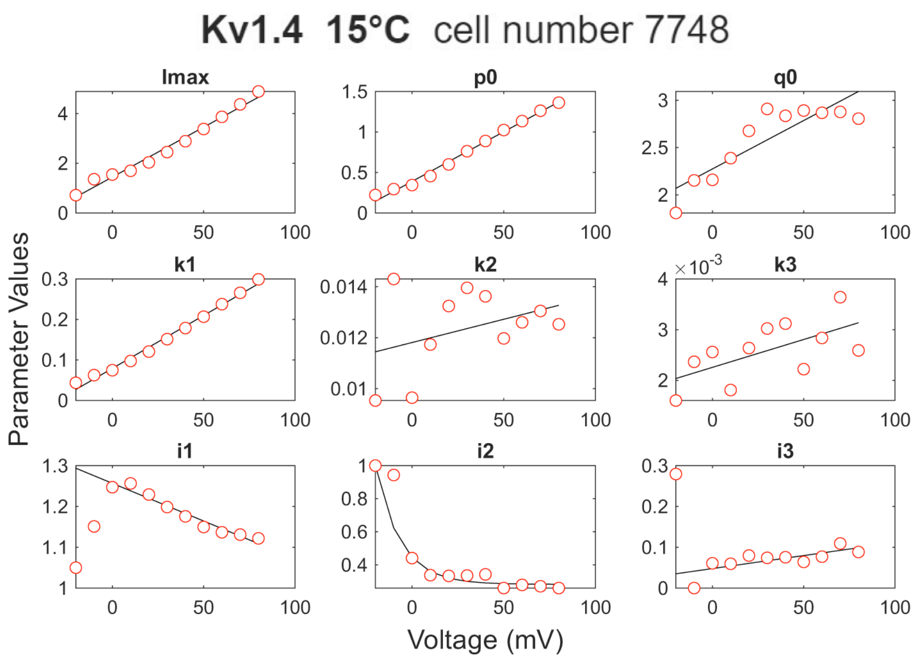


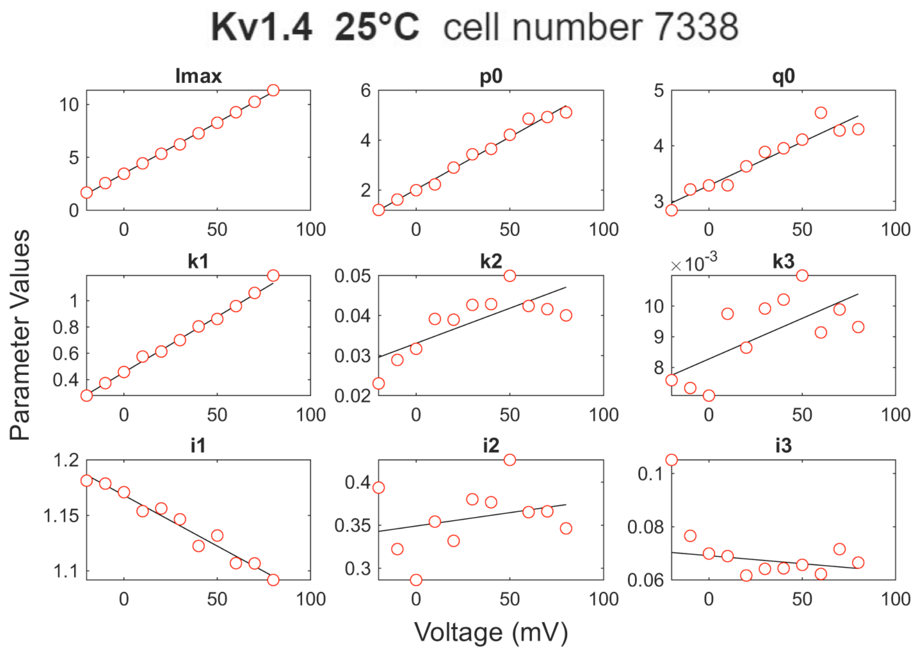


*Figure S4 Continued on next page*

*Figure S4 continued*

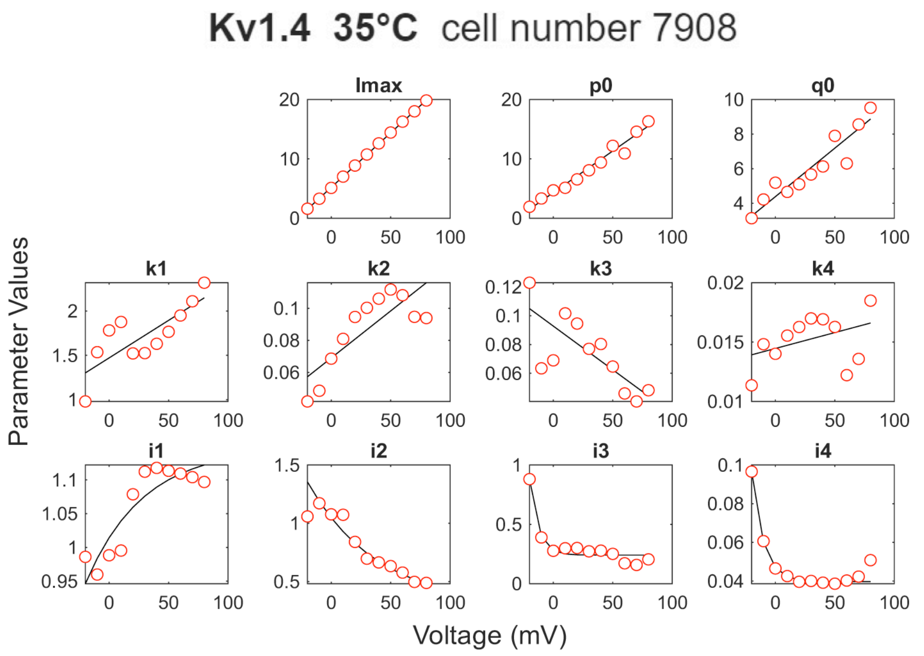


**Figure S4**. MS model parameters for fitting Kv1.4 activation currents at three temperatures. Red circles represent the parameter values obtained from the fitting. They are further fitted by linear functions (black lines) except that $i_2$ of Kv1.4 (15 ºC), and $i_1$, $i_2$, $i_3$, and $i_4$ of Kv1.4 (35 ºC) are fitted by exponential functions.

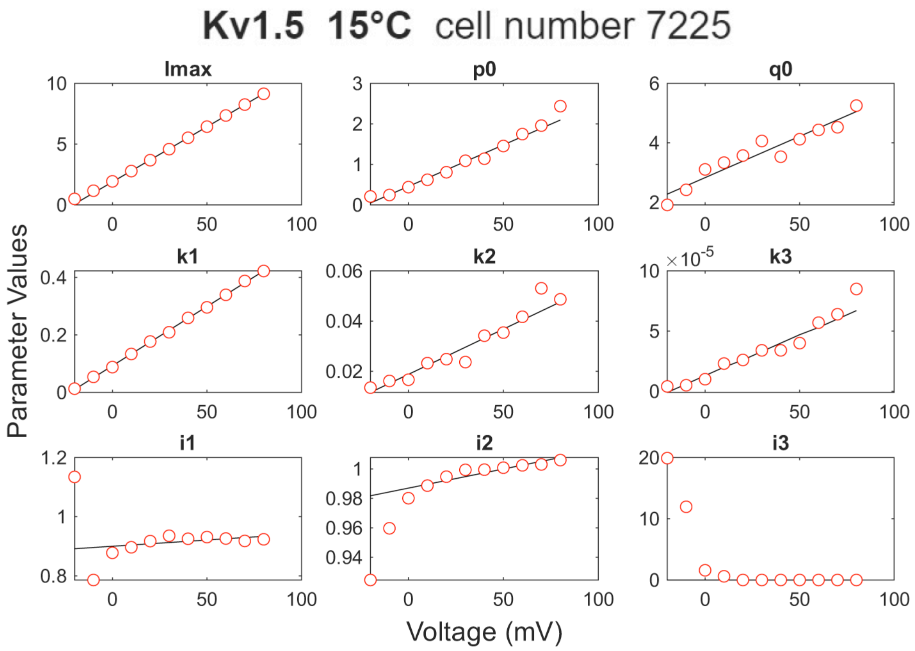


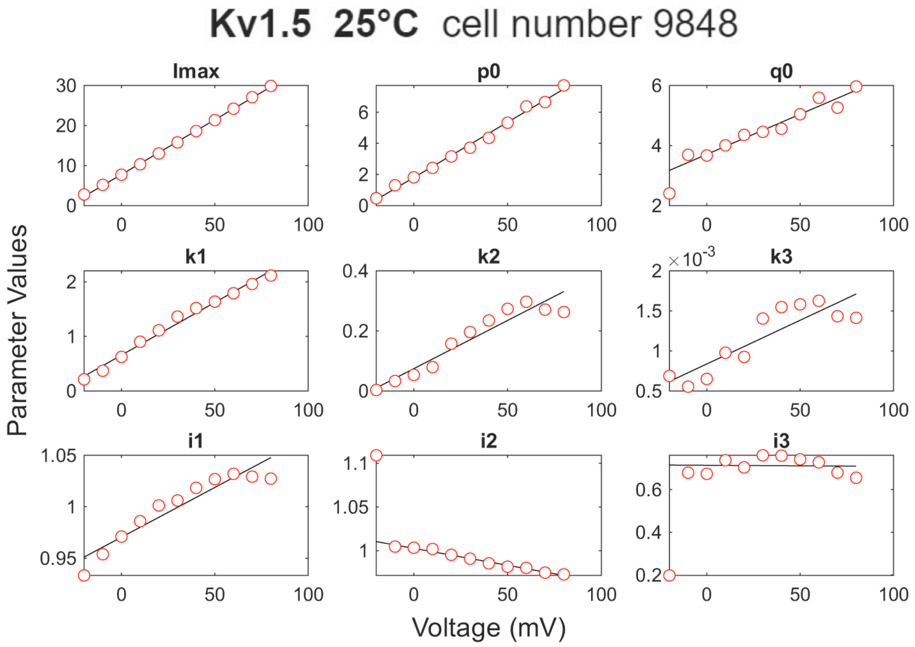


*Figure S5 Continued on next page*

*Figure S5 continued*

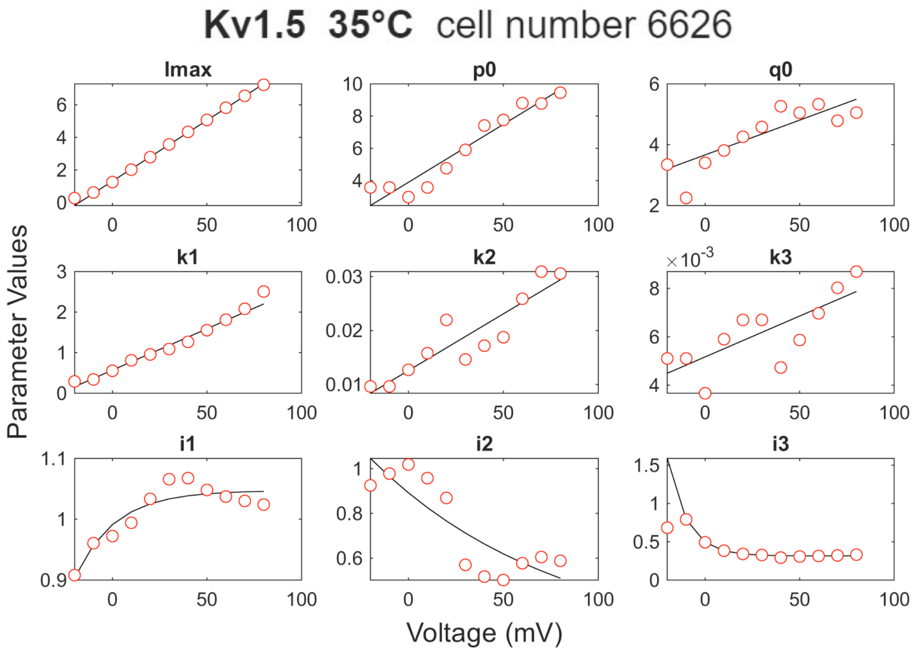


**Figure S5**. MS model parameters for fitting Kv1.5 activation currents at three temperatures. Red circles represent the parameter values obtained from the fitting, which are further fitted by linear functions (black lines) except the Kv1.5 (35 ºC) results where $i_1$, $i_2$, and $i_3$ are fitted by exponential functions.

**Kv1.6 15°C** cell number 7728

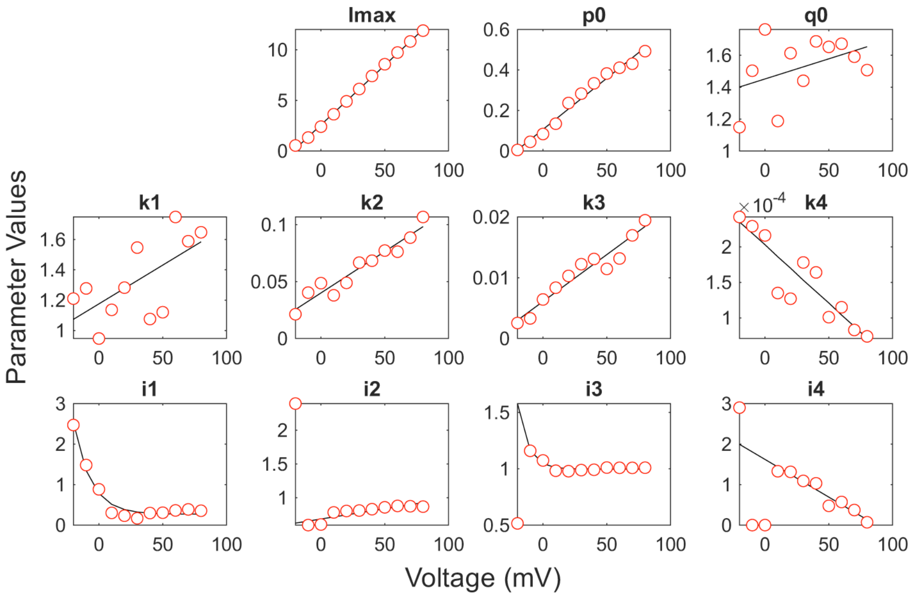


**Kv1.6 25°C** cell number 9946

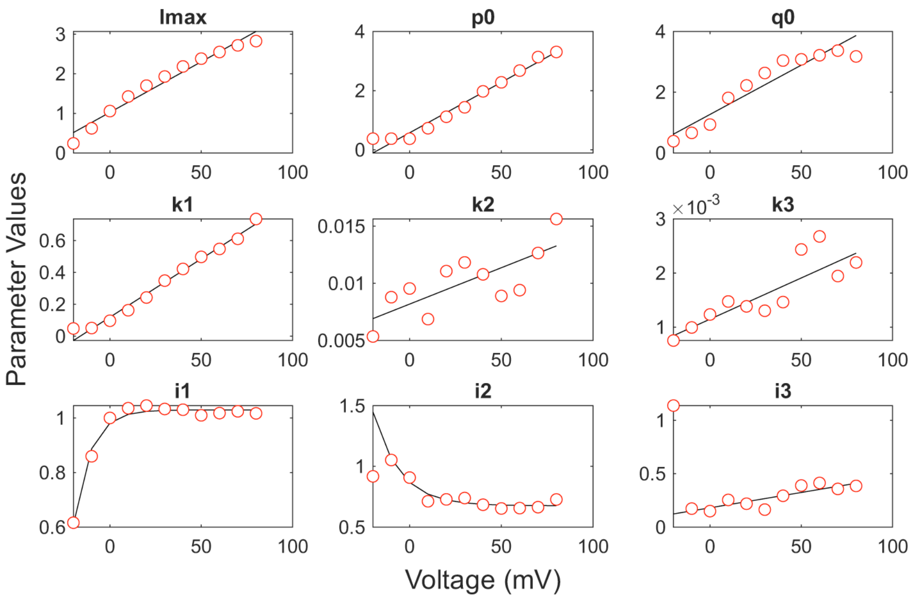


*Figure S6 Continued on next page*

*Figure S6 continued*

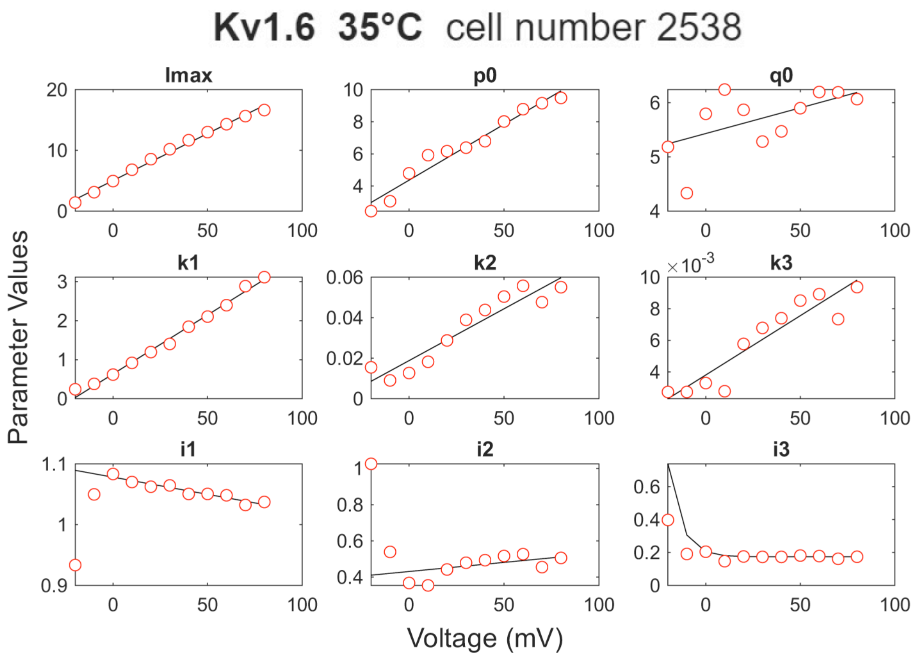


**Figure S6**. MS model parameters for fitting Kv1.6 activation currents at three temperatures. Red circles represent the parameter values obtained from the fitting. They are further fitted by linear functions (black lines) except that $i_1$ and $i_3$ of Kv1.6 (15 ºC), $i_1$ and $i_2$ of Kv1.6 (25 ºC), and $i_3$ of Kv1.6 (35 ºC) are fitted by exponential functions.

**Kv1.7 15°C cell number 12427**

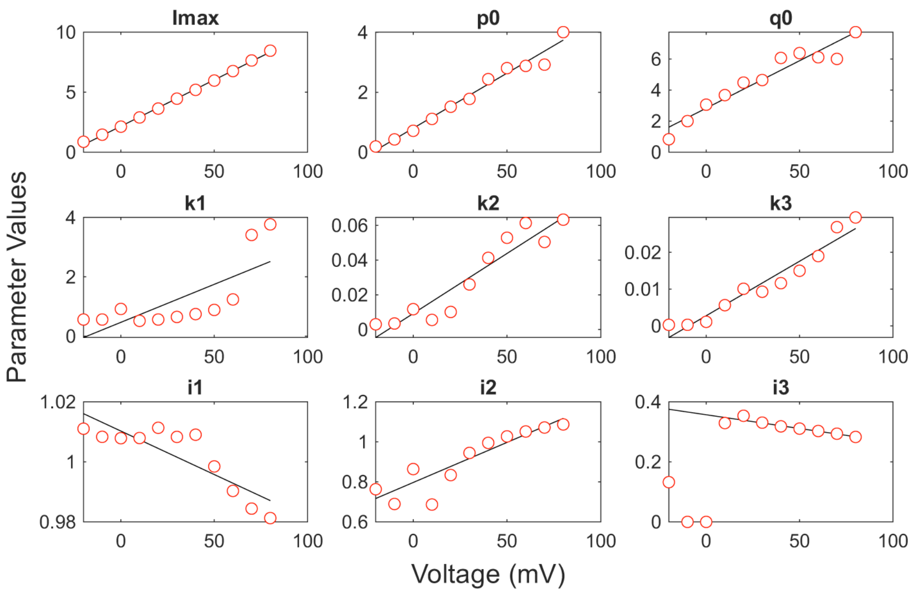


**Kv1.7 25°C cell number 19551**

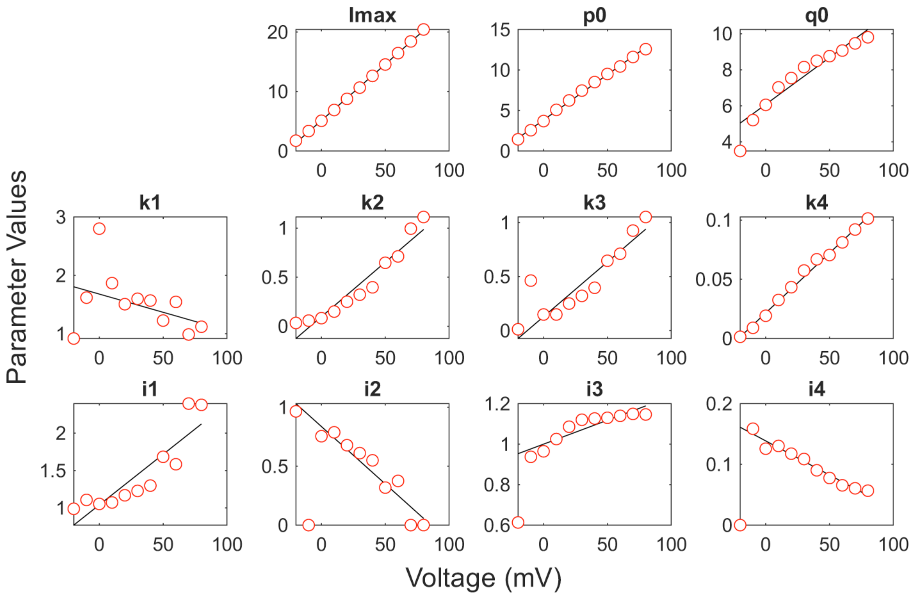


*Figure S7 Continued on next page*

*Figure S7 continued*

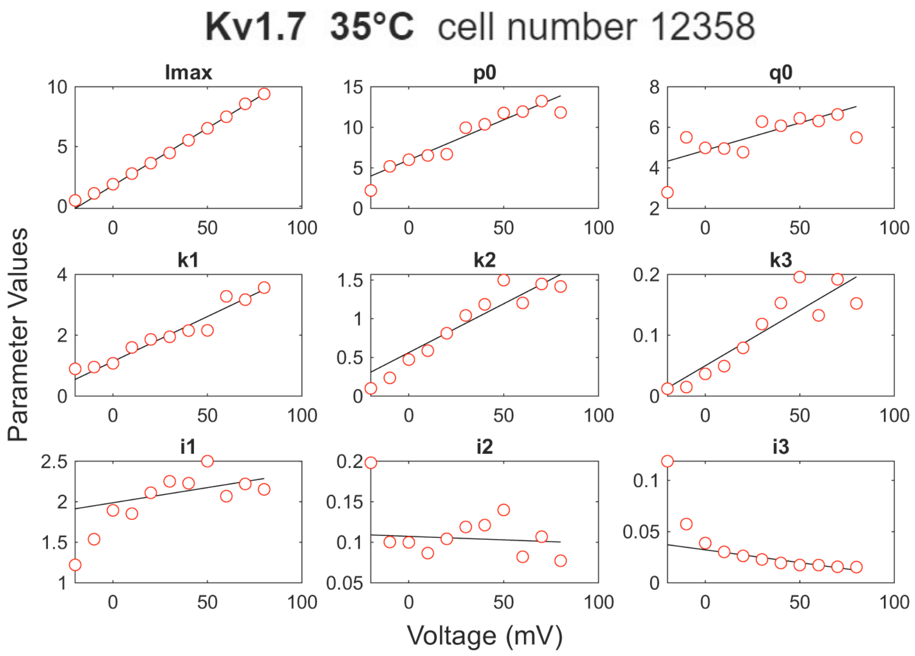


**Figure S7**. MS model parameters for fitting Kv1.7 activation currents at three temperatures. Red circles represent the parameter values obtained from the fitting, which are further fitted by linear functions (black lines).

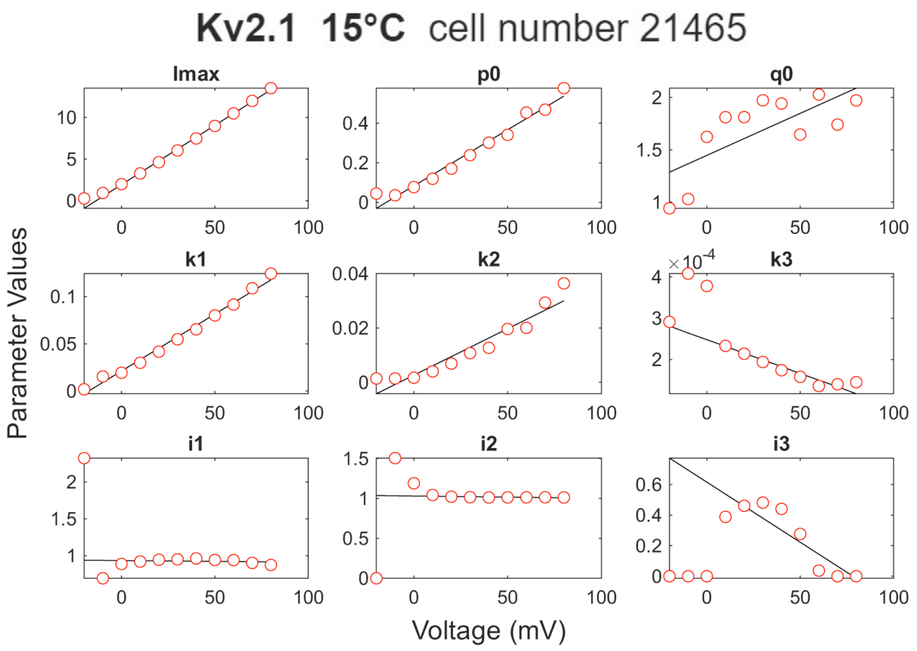


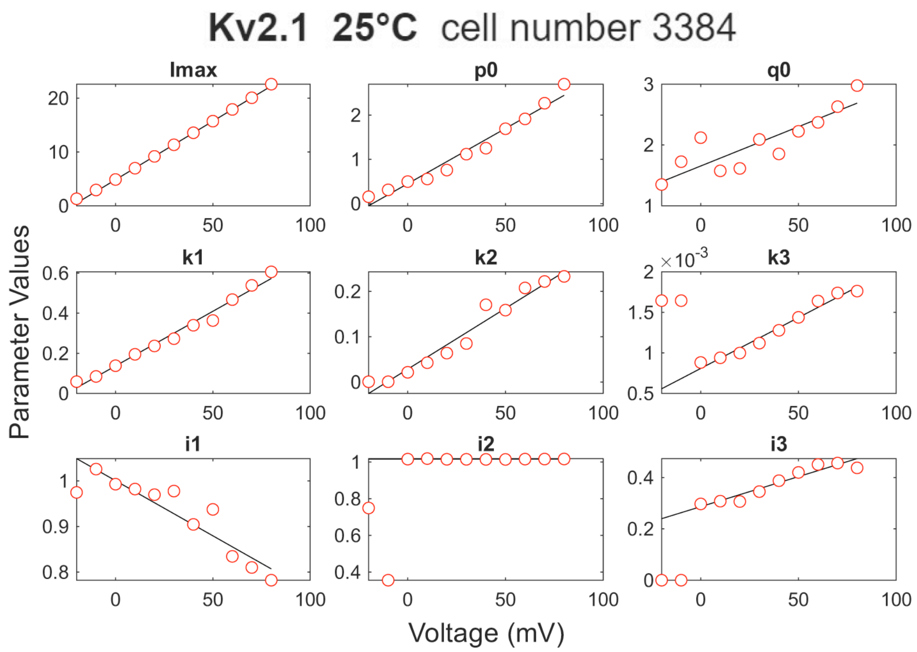


*Figure S8 Continued on next page*

*Figure S8 continued*

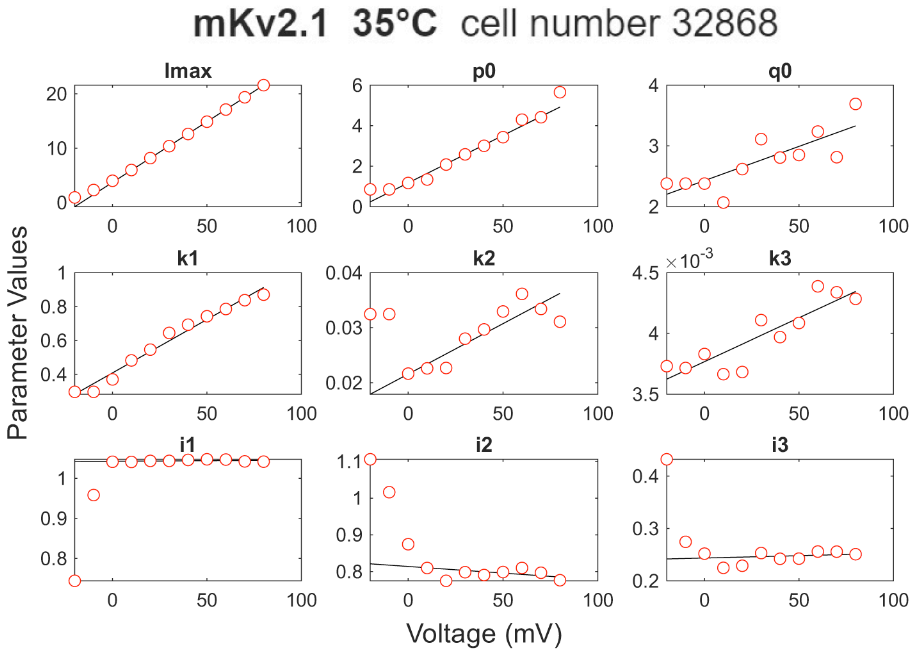


**Figure S8**. MS model parameters for fitting Kv2.1 activation currents at three temperatures. Red circles represent the parameter values obtained from the fitting, which are further fitted by linear functions (black lines).

### Kv2.2 15°C cell number 3190

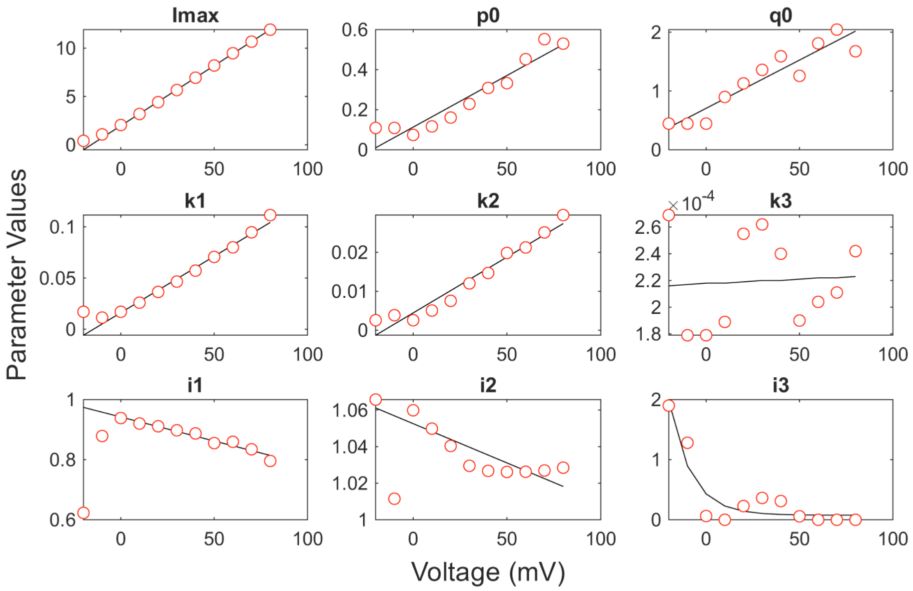


### Kv2.2 25°C cell number 9932

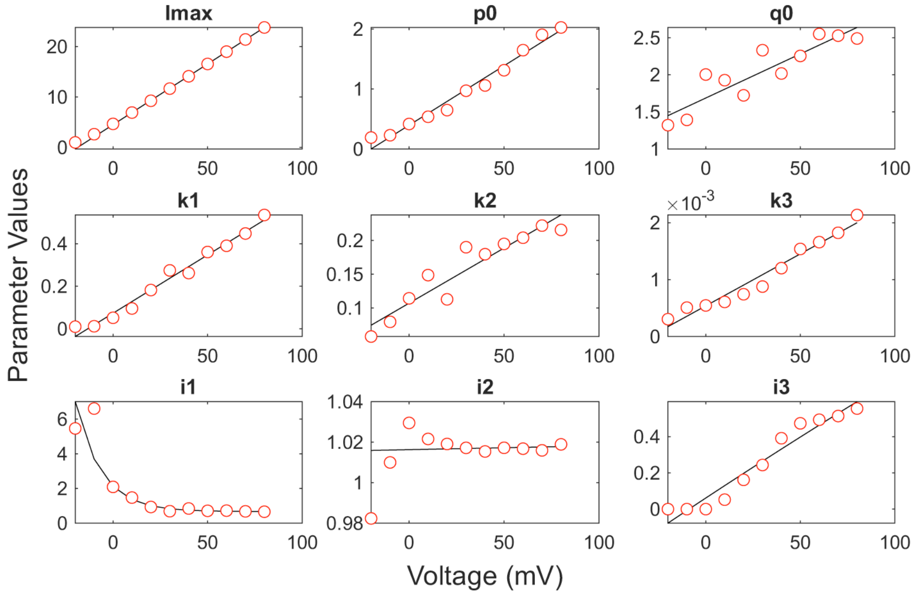


*Figure S9 Continued on next page*

*Figure S9 continued*

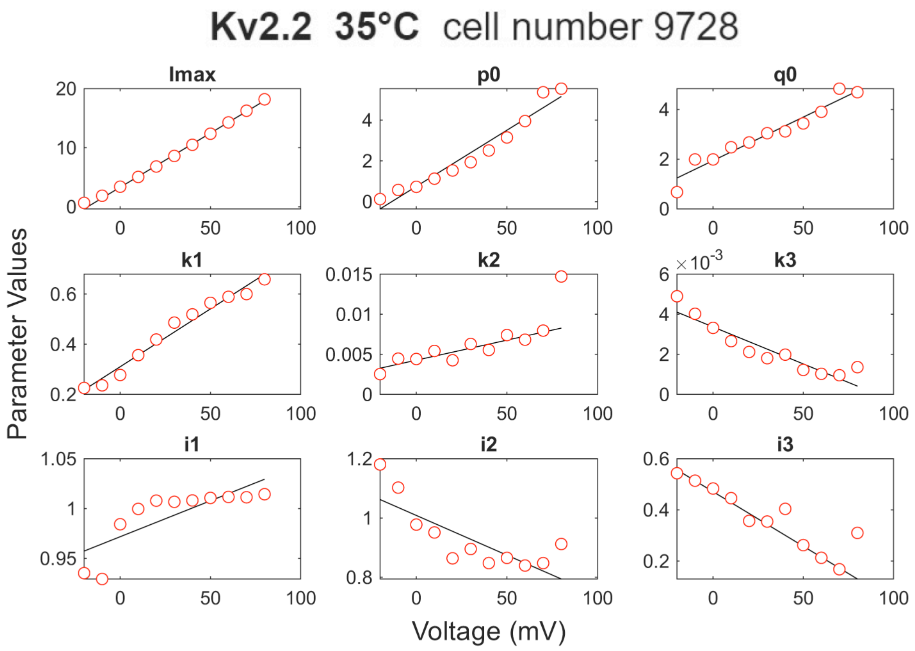


**Figure S9**. MS model parameters for fitting Kv2.2 activation currents at three temperatures. Red circles represent the parameter values obtained from the fitting. They are further fitted by linear functions (black lines) except that $i_3$ of Kv2.2 (15 ºC) and $i_1$ of Kv2.2 (25 ºC) are fitted by exponential functions.

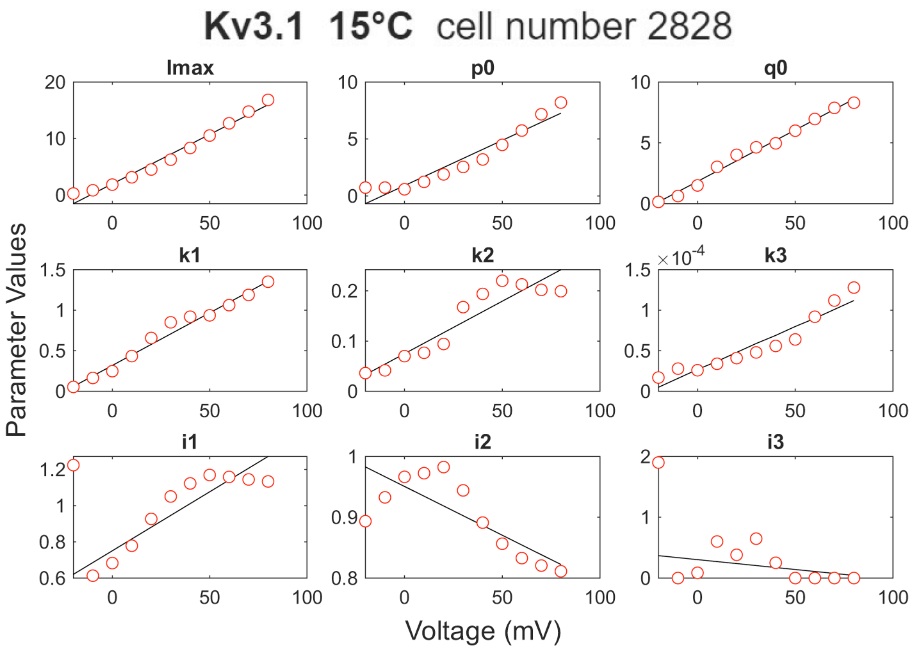


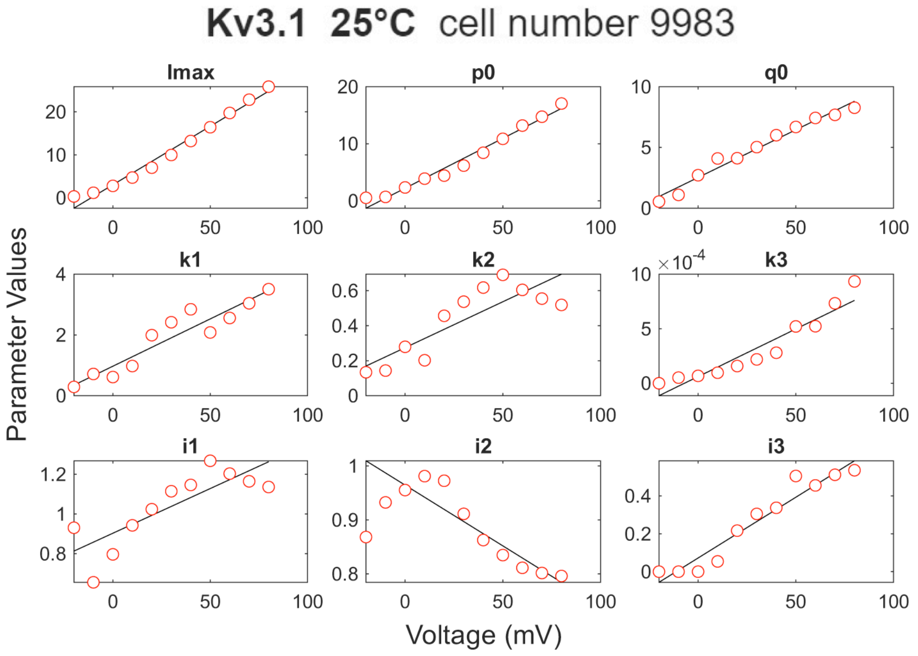


*Figure S10 Continued on next page*

*Figure S10 continued*

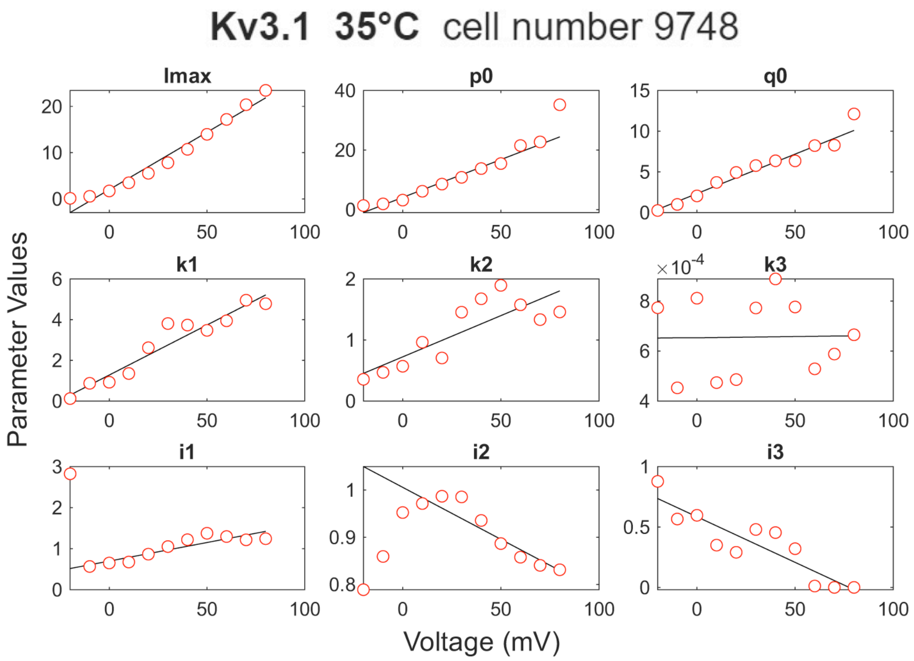


**Figure S10**. MS model parameters for fitting Kv3.1 activation currents at three temperatures. Red circles represent the parameter values obtained from the fitting, which are further fitted by linear functions (black lines).

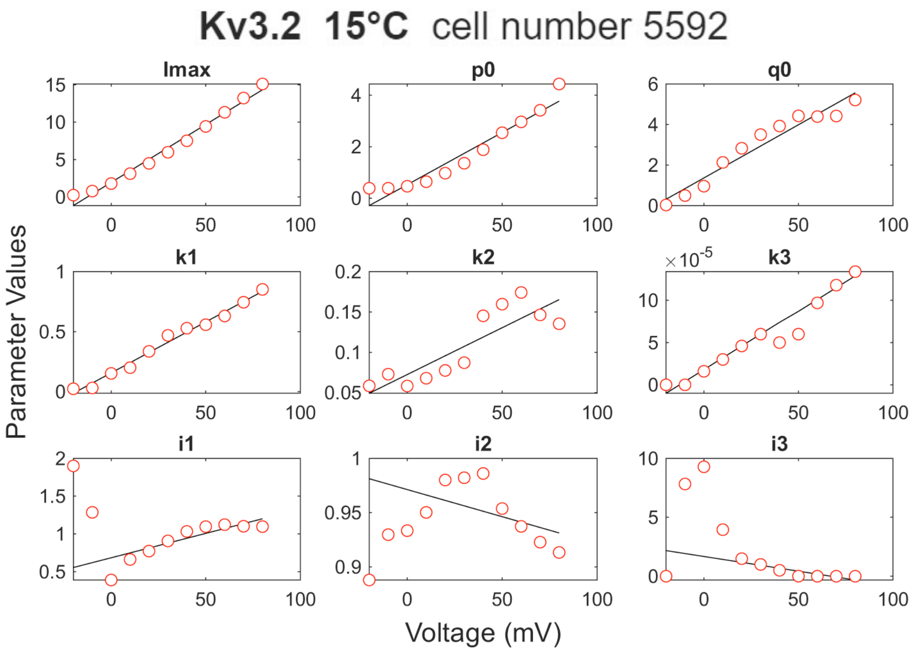


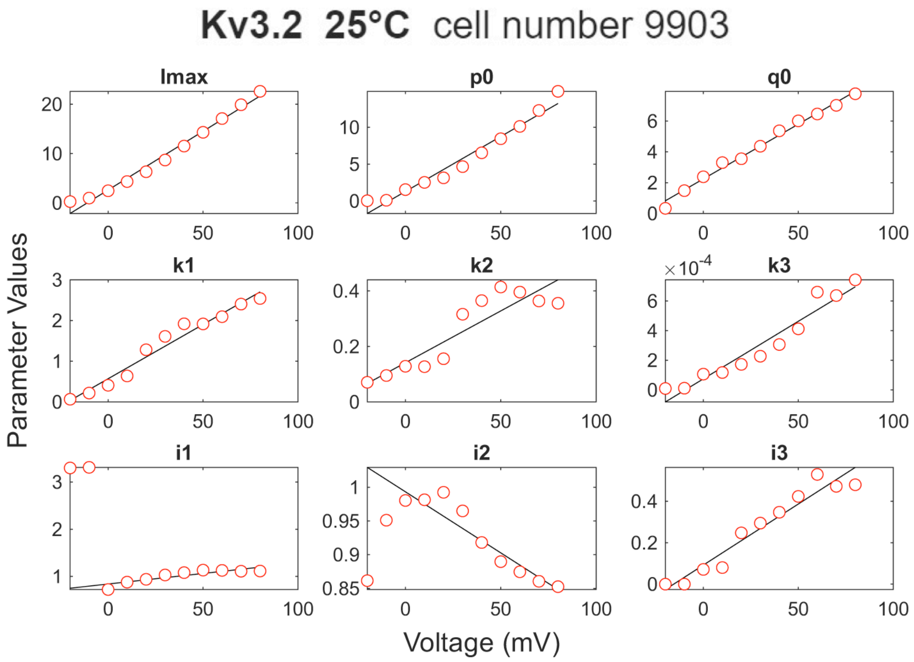


*Figure S11 Continued on next page*

*Figure S11 continued*

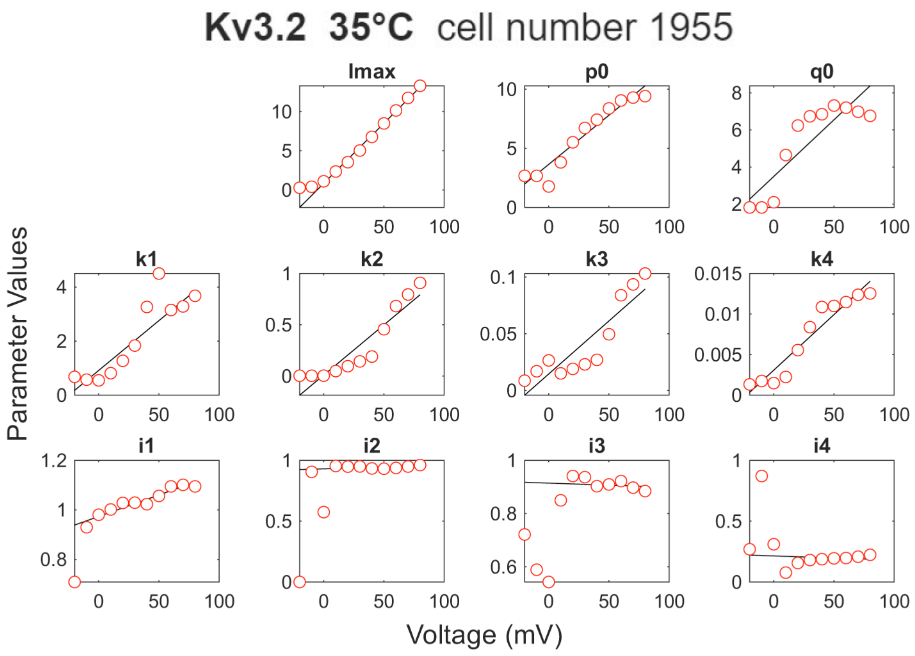


**Figure S11**. MS model parameters for fitting Kv3.2 activation currents at three temperatures. Red circles represent the parameter values obtained from the fitting, which are further fitted by linear functions (black lines).

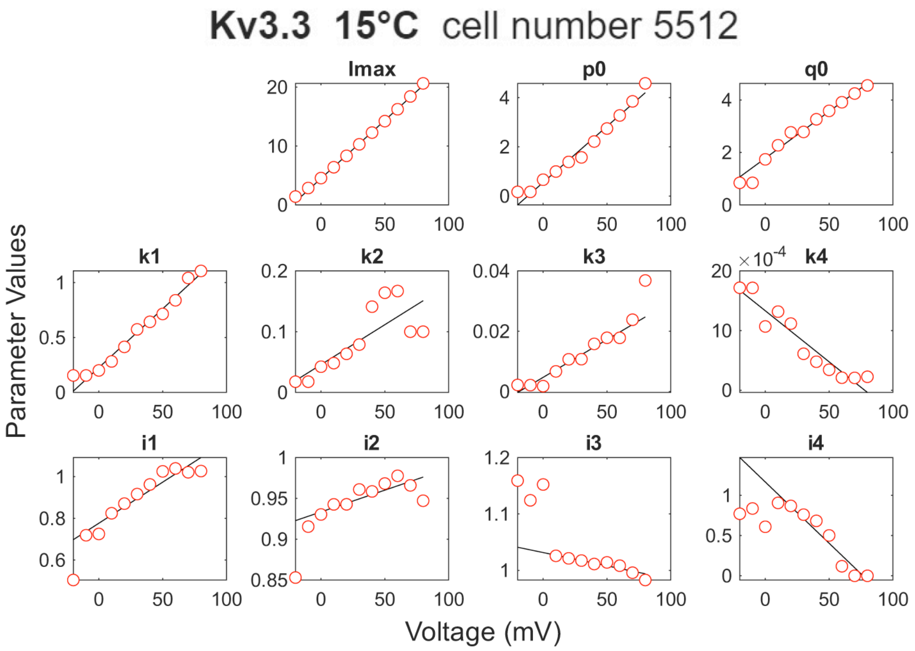


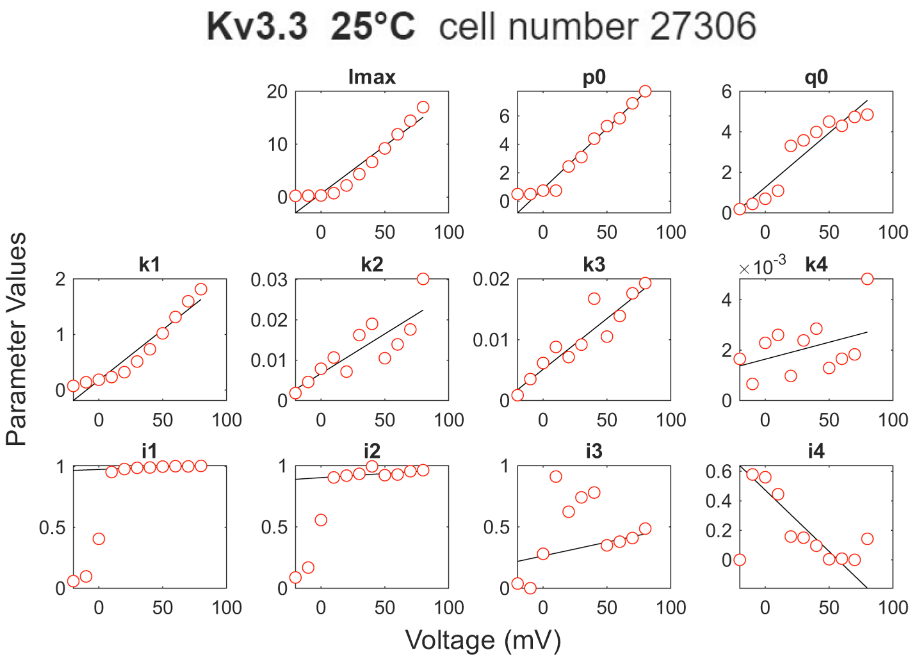


*Figure S12 Continued on next page*

*Figure S12 continued*

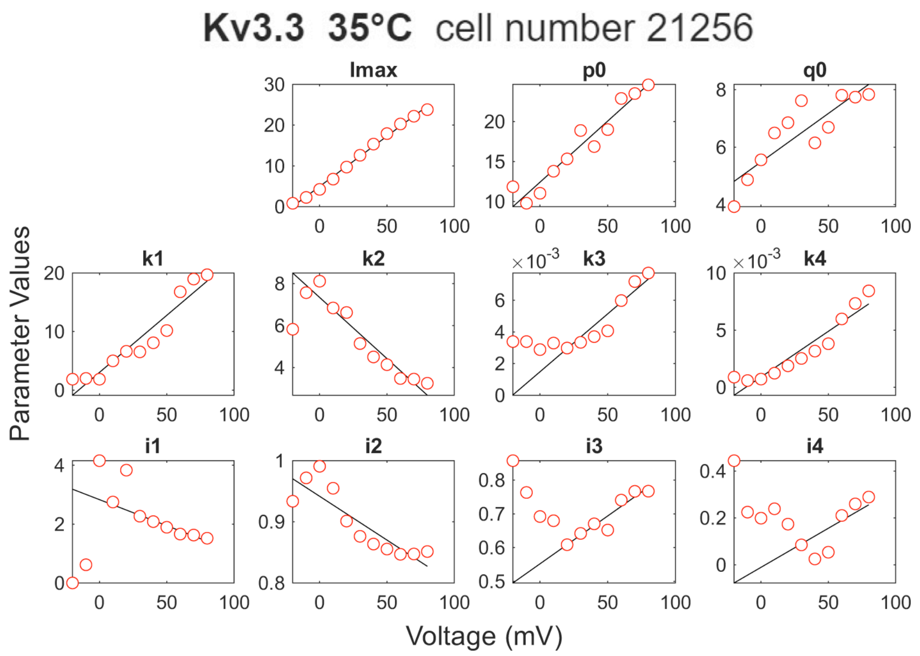


**Figure S12**. MS model parameters for fitting Kv3.3 activation currents at three temperatures. Red circles represent the parameter values obtained from the fitting, which are further fitted by linear functions (black lines).

### Kv3.4 15°C cell number 5561

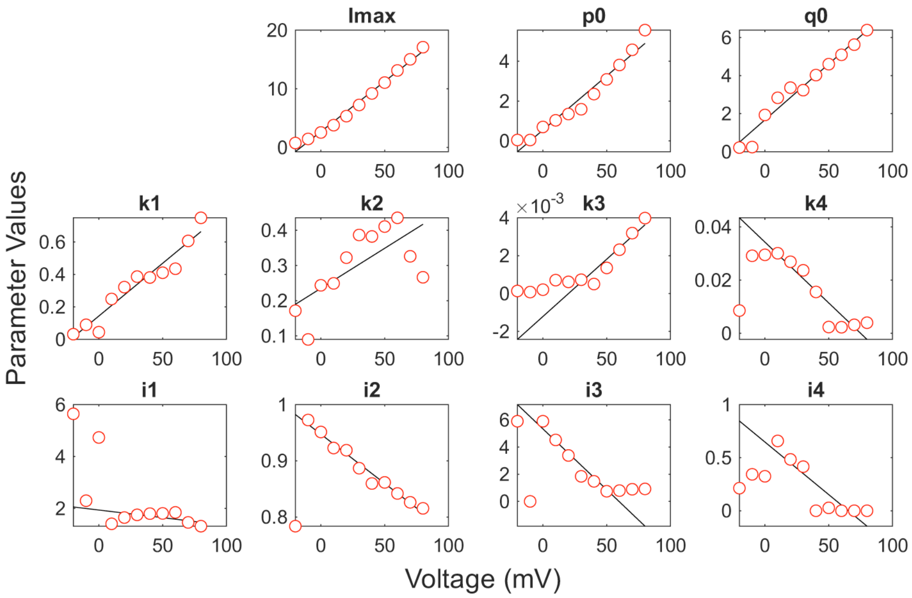


### Kv3.4 25°C cell number 12618

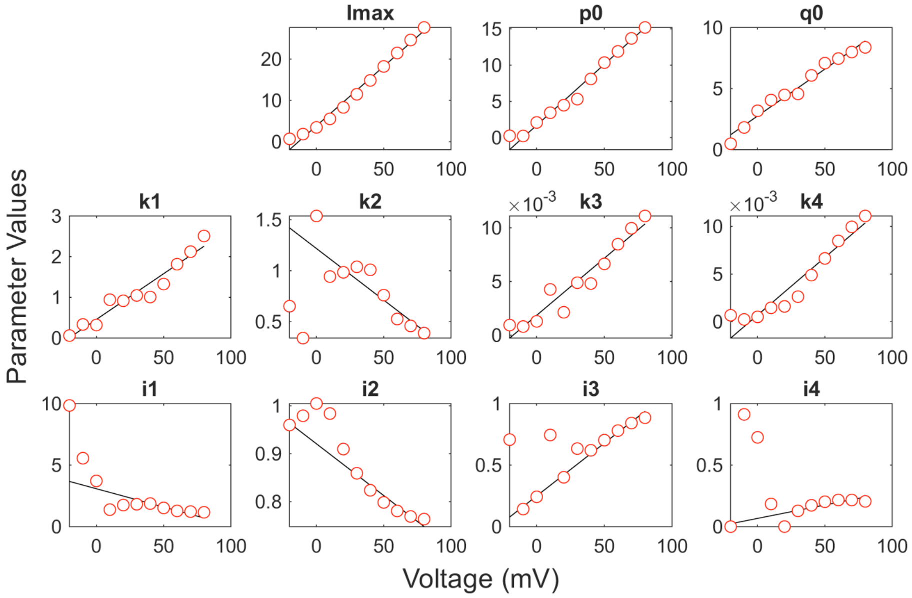


*Figure S13 Continued on next page*

*Figure S13 continued*

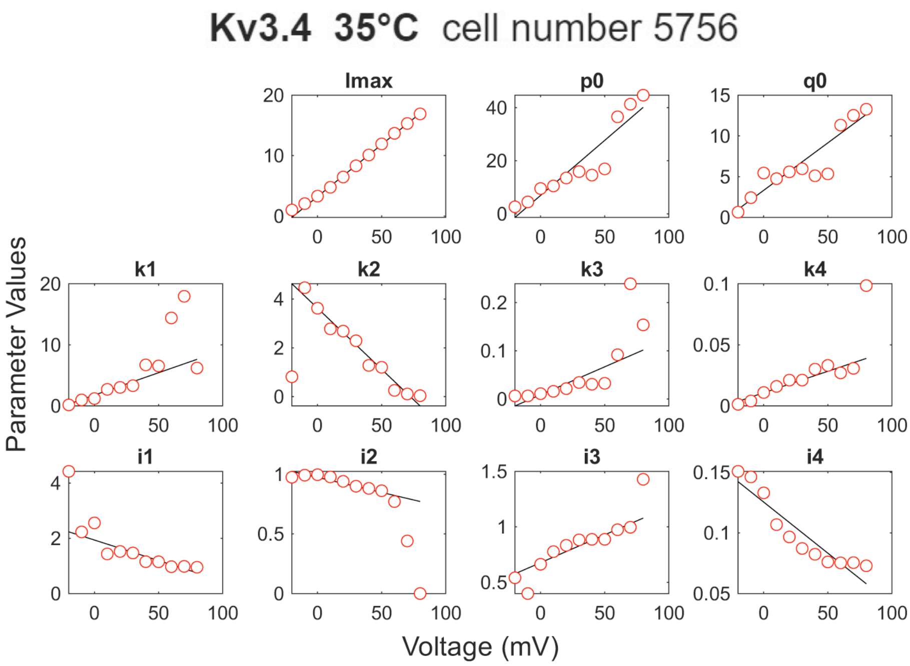


**Figure S13**. MS model parameters for fitting Kv3.4 activation currents at three temperatures. Red circles represent the parameter values obtained from the fitting, which are further fitted by linear functions (black lines).

### Kv4.1 15°C cell number 8001

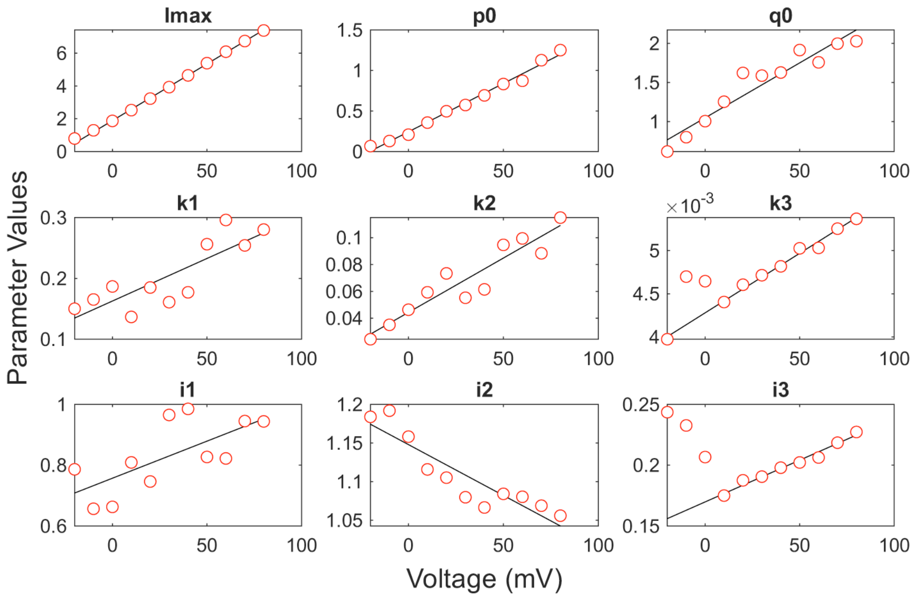


### Kv4.1 25°C cell number 16370

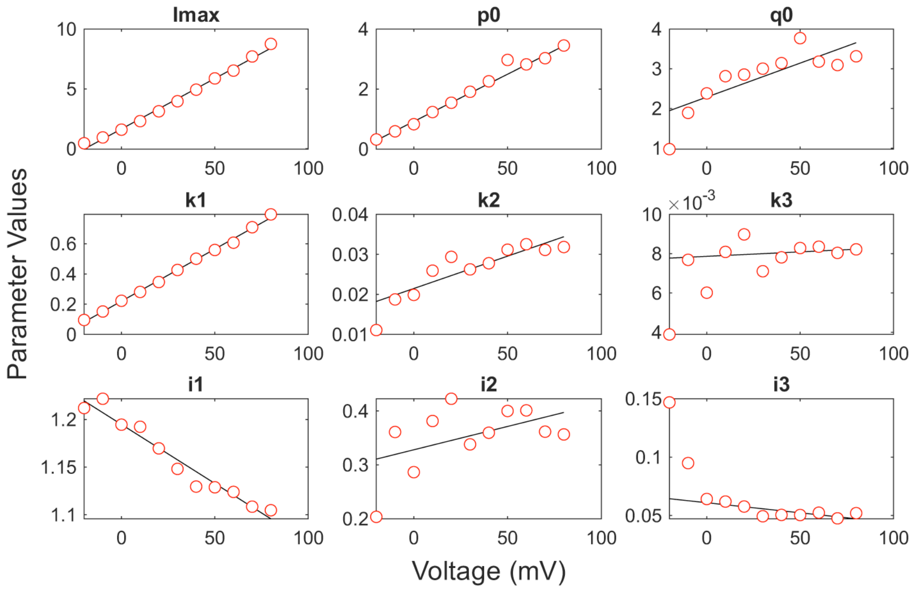


*Figure S14 Continued on next page*

*Figure S14 continued*

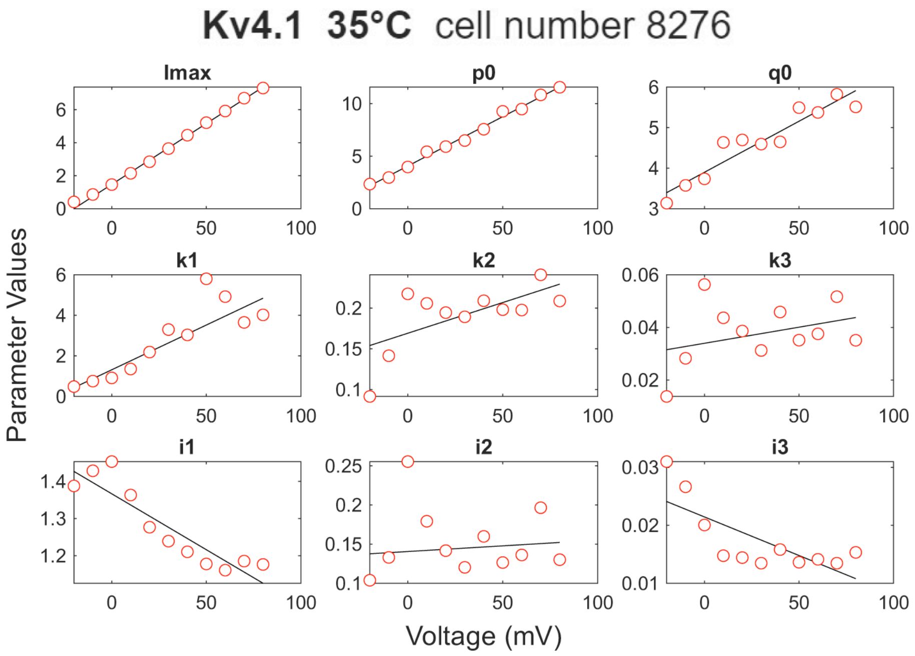


**Figure S14**. MS model parameters for fitting Kv4.1 activation currents at three temperatures. Red circles represent the parameter values obtained from the fitting, which are further fitted by linear functions (black lines).

### Kv4.2 15°C cell number 6496

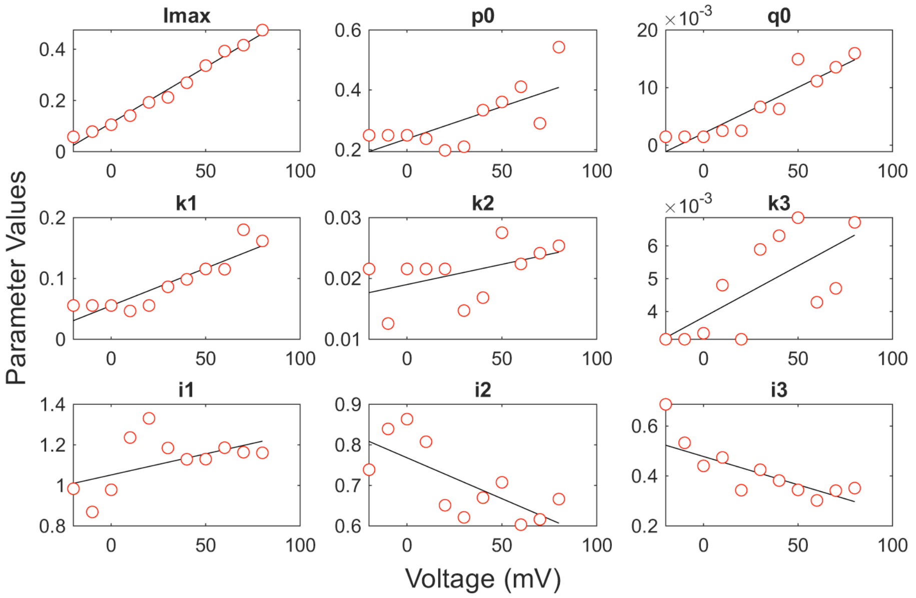


### Kv4.2 25°C cell number 8309

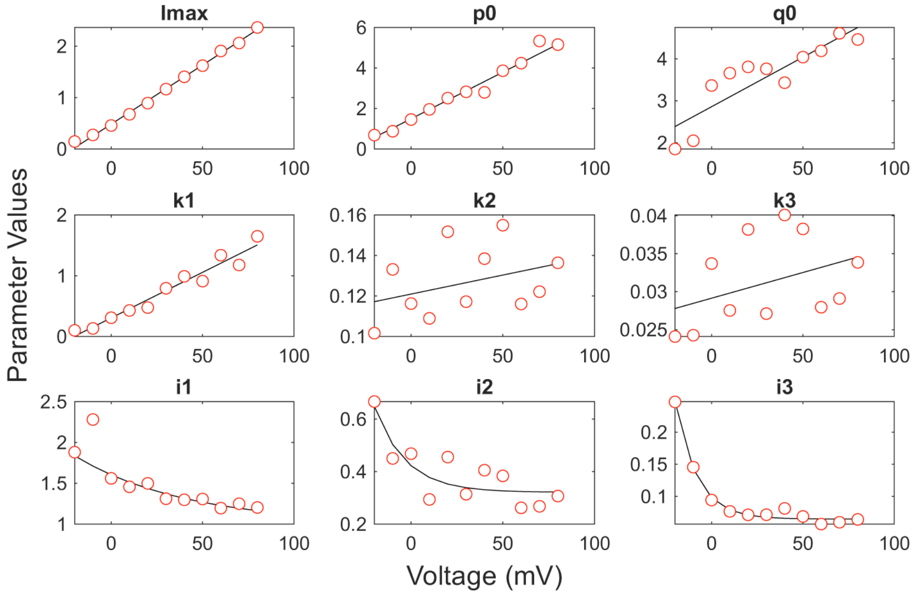


*Figure S15 Continued on next page*

*Figure S15 continued*

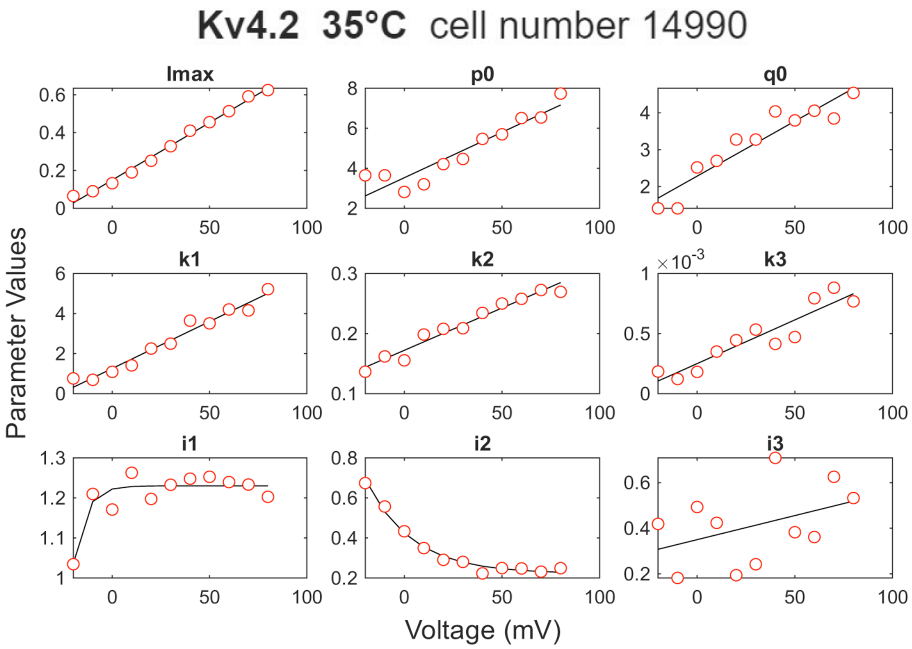


**Figure S15**. MS model parameters for fitting Kv4.2 activation currents at three temperatures. Red circles represent the parameter values obtained from the fitting. They are further fitted by linear functions (black lines) except that $i_1$, $i_2$, and $i_3$ of Kv4.2 (25 ºC), and $i_1$ and $i_2$ of Kv4.2 (35 ºC) are fitted by exponential functions.

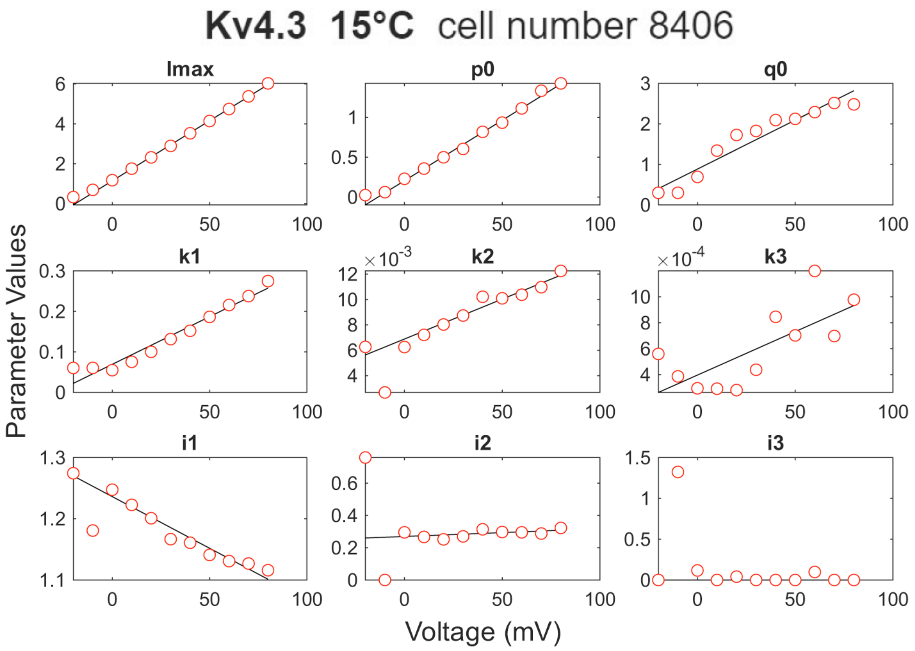


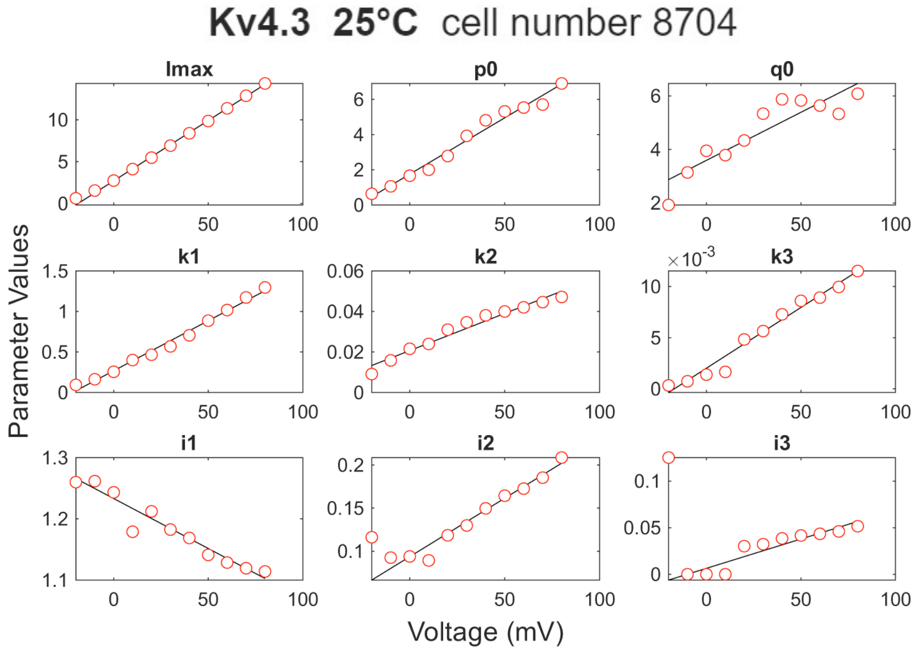


*Figure S16 Continued on next page*

*Figure S16 continued*

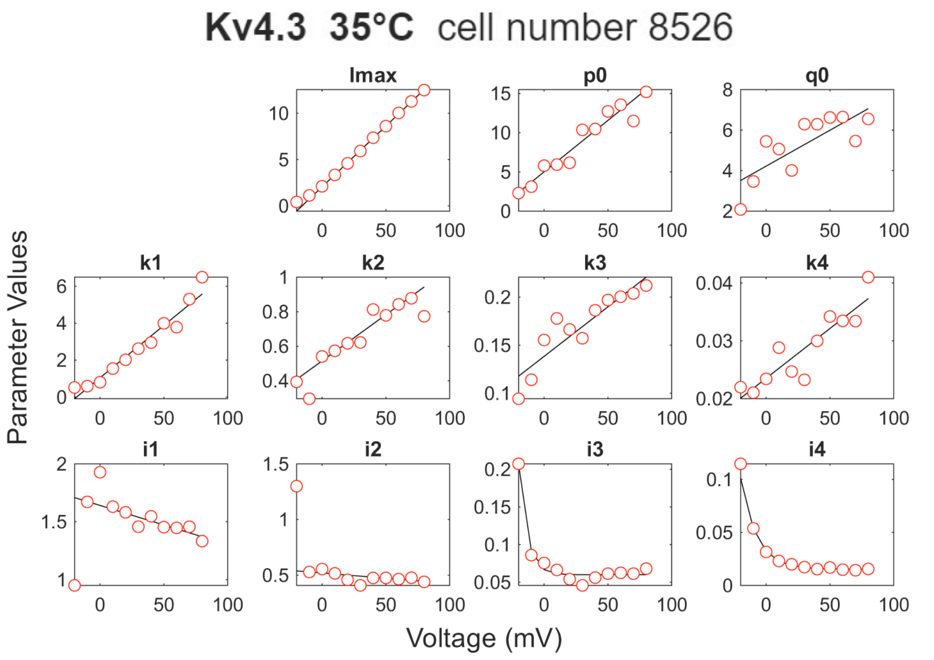


**Figure S16**. MS model parameters for fitting Kv4.3 activation currents at three temperatures. Red circles represent the parameter values obtained from the fitting, which are further fitted by linear functions (black lines) except the Kv4.3 (35 ºC) results where $i_3$ and $i_4$ are fitted by exponential functions.

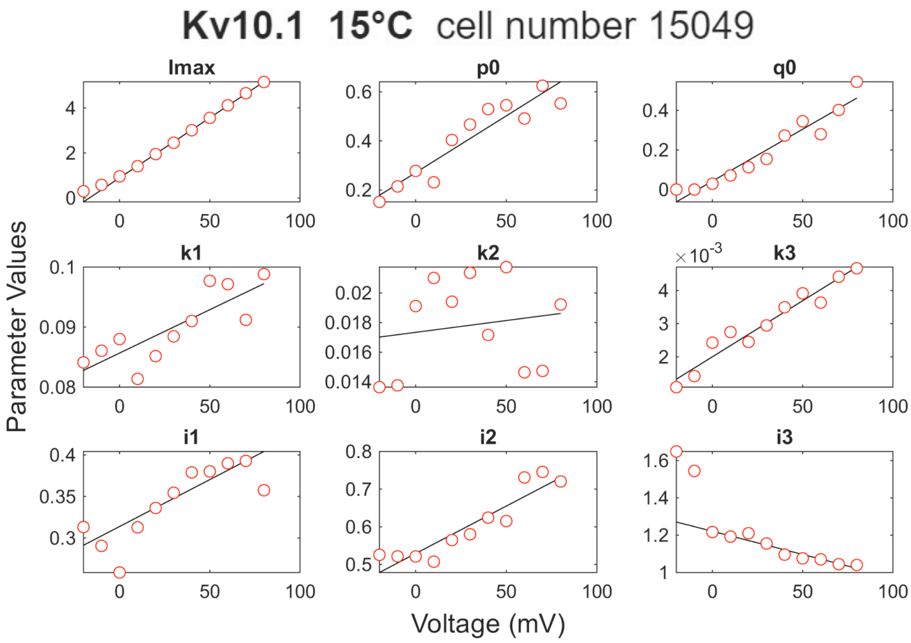


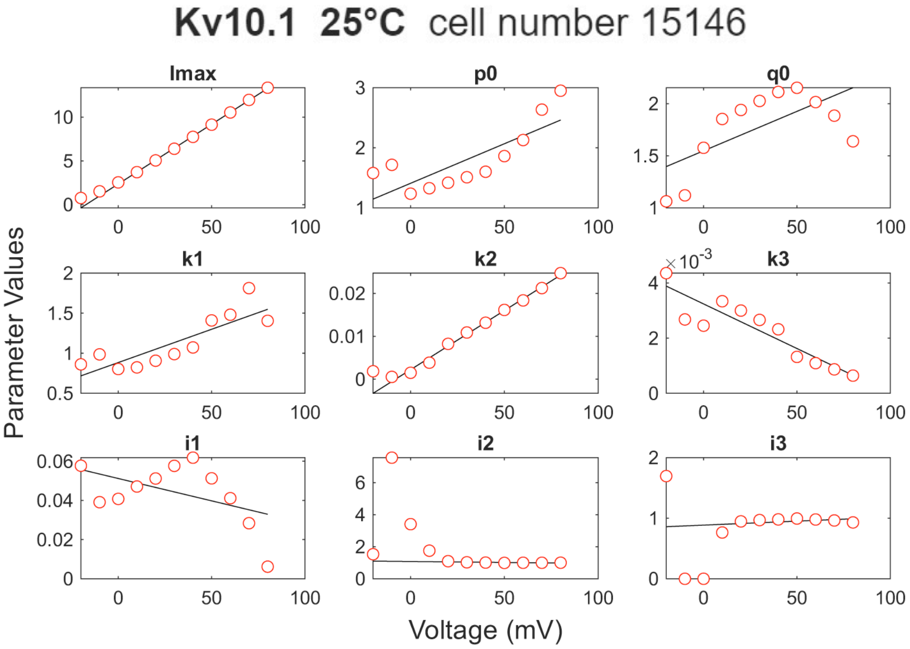


*Figure S17 Continued on next page*

*Figure S17 continued*

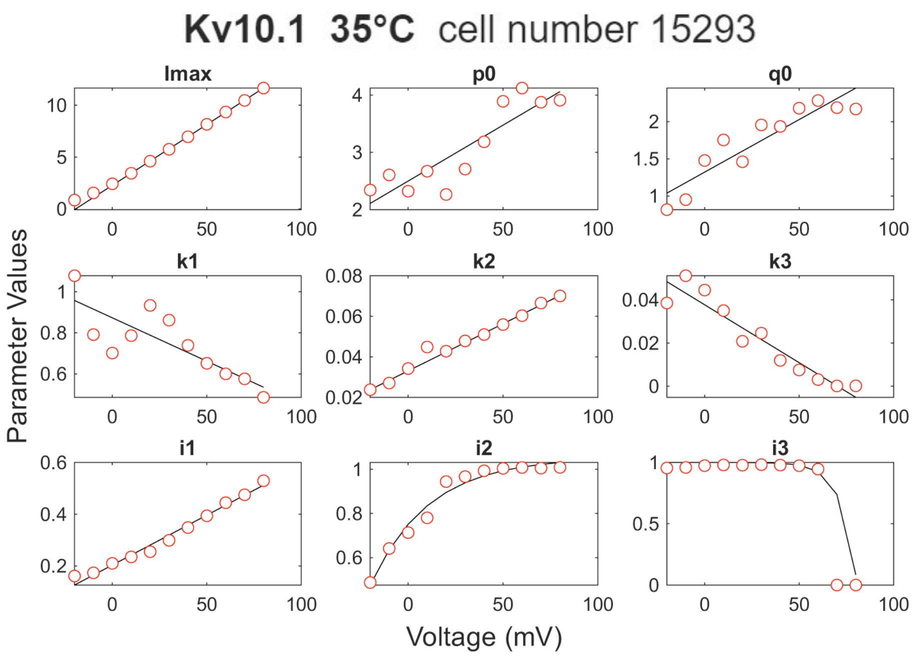


**Figure S17**. MS model parameters for fitting Kv10.1 activation currents at three temperatures. Red circles represent the parameter values obtained from the fitting, which are further fitted by linear functions (black lines) except the Kv10.1 (35 ºC) results where $i_2$ and $i_3$ are fitted by exponential functions.

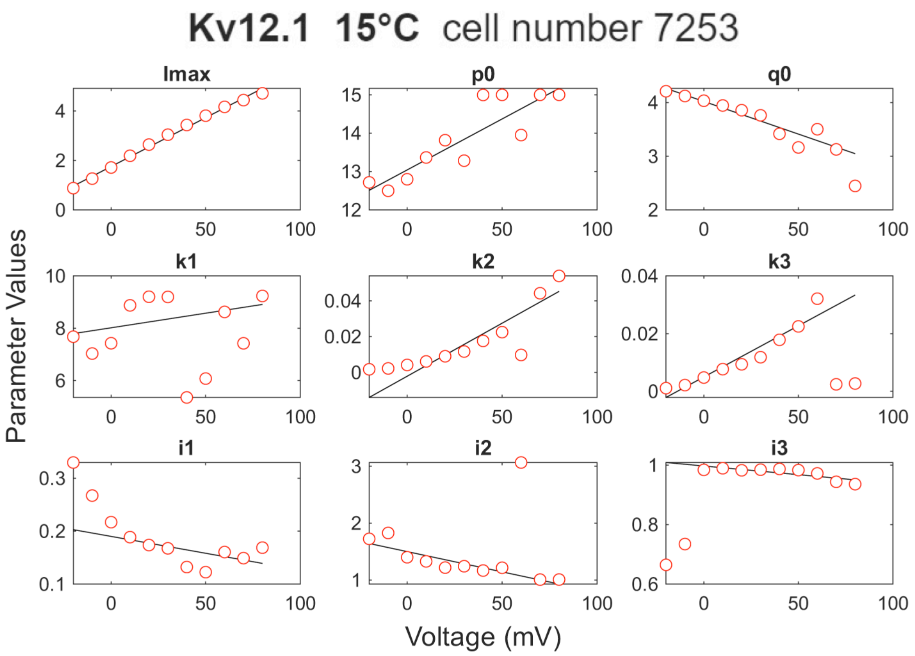


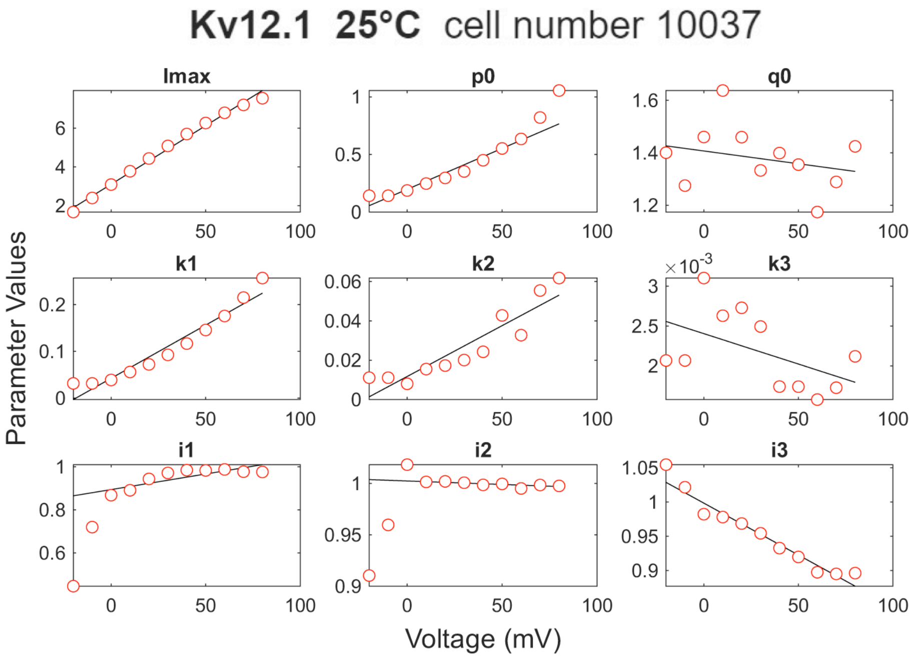


*Figure S18 Continued on next page*

*Figure S18 continued*

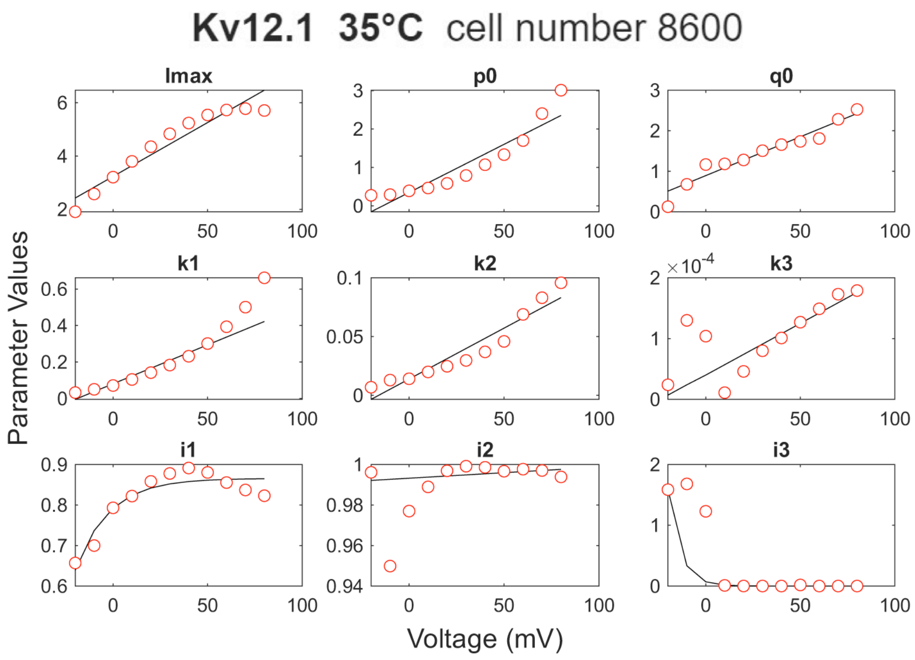


**Figure S18**. MS model parameters for fitting Kv12.1 activation currents at three temperatures. Red circles represent the parameter values obtained from the fitting, which are further fitted by linear functions (black lines) except the Kv12.1 (35 ºC) results where $i_1$ and $i_3$ are fitted by exponential functions.
